# Natural Killer Cell Dysfunction In Human Bladder Cancer Is Caused By Tissue-Specific Suppression of SLAMF6 Signaling

**DOI:** 10.1101/2024.04.30.591366

**Authors:** Adam M. Farkas, Dina Youssef, Michelle A. Tran, Sreekumar Balan, Jenna H. Newman, François Audenet, Harry Anastos, Leandra G. Velazquez, Ante Peros, Aparna Ananthanarayanan, Jorge Daza, Elena Gonzalez-Gugel, Keerthi Sadanala, Jakob Theorell, Matthew D. Galsky, Amir Horowitz, John P. Sfakianos, Nina Bhardwaj

## Abstract

NK cells are innate lymphocytes critical for surveillance of viruses and tumors, however the mechanisms underlying NK cell dysfunction in cancer are incompletely understood. We assessed the effector function of NK cells from bladder cancer patients and found severe dysfunction in NK cells derived from tumors versus peripheral blood. While both peripheral and tumor-infiltrating NK cells exhibited conserved patterns of inhibitory receptor over-expression, this did not explain the observed defects in NK surveillance in bladder tumors. Rather, TME-specific TGF-β and metabolic perturbations such as hypoxia directly suppressed NK cell function. Specifically, an oxygen-dependent reduction in signaling through SLAMF6 was mechanistically responsible for poor NK cell function, as tumor-infiltrating NK cells cultured *ex vivo* under normoxic conditions exhibited complete restoration of function, while deletion of SLAMF6 abrogated NK cell cytolytic function even under normoxic conditions. Collectively, this work highlights the role of tissue-specific factors in dictating NK cell function, and implicates SLAMF6 signaling as a rational target for immuno-modulation to improve NK cell function in bladder cancer.

**Graphical Abstract:** 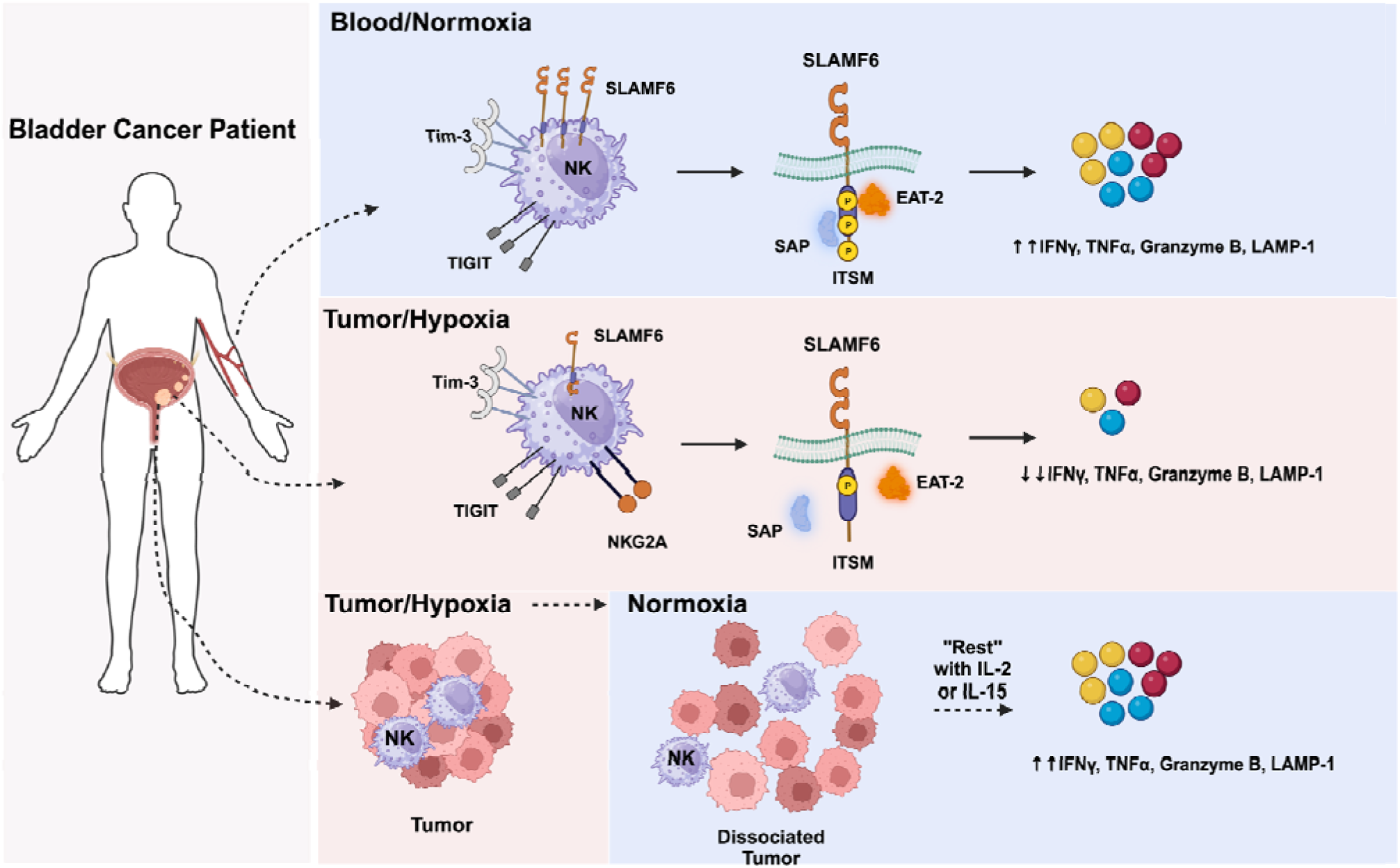

## Introduction

Urothelial cell carcinoma, or bladder cancer (BCa) is the 6^th^ most commonly diagnosed cancer worldwide, resulting in 150,000 deaths annually^1^. Clinically, BCa is classified as non-muscle invasive (NMI), muscle-invasive (MI), or metastatic, with each stage having different prognoses and treatment. Patients with high-risk NMI are generally resected then treated with intravesical BCG, a live attenuated preparation of *Mycobacterium bovis*, highlighting the amenability of bladder tumors to respond to immunotherapy. More recently, subsets of patients with metastatic and locally-advanced disease have benefited from T cell-centric, immune checkpoint blockade (ICB) using monoclonal antibodies (mAb) interfering with the PD-1: PD-L1 axis. Despite this progress, the average response rate for ICB is ∼20-30% ^2–7^, highlighting the need to identify additional targets that improve response rates. Natural Killer cells (NK) are innate lymphocytes critical for host defense against viral pathogens and transformed cells ^8–10^. Unlike T and B cells, NK cells lack a somatically recombined, antigen-specific receptor. Instead, targets are generally recognized as cells that have down-regulated Class I HLA, and up-regulated molecules associated with cellular stress ^11–13^. NK cell function is further tuned by integrating signals transmitted by a diverse array of activating and inhibitory receptors ^13,14^. NK cell effector functions include the production of inflammatory cytokines such as IFNγ and TNFα, recruitment and maintenance of dendritic cells via production of CCL5, GM-CSF, FLT3-L and XCL1/2, antibody-dependent cellular cytotoxicity (ADCC), and direct killing via release of cytolytic granules containing perforin and granzymes^15^ . The two major subsets of human NK cells differ in effector function, with less mature CD56^bright^CD16^-^ NK cells responsible for most cytokine production, and more mature CD56^dim^CD16^+^ cells having enhanced cytotoxic potential^16,17^.

During cancer and chronic viral infection, T cells acquire a phenotype of immune exhaustion marked by upregulation of inhibitory receptors including PD-1, CTLA-4, Tim-3 and TIGIT, and defined by effector dysfunction^18^. However, whether a parallel process affects NK cells remains incompletely understood. Additionally, there is little insight pertaining to differences in NK cell phenotype between primary tumor and distal sites like peripheral blood. We have previously shown that NK cells from peripheral blood (pbNK) up-regulate the inhibitory receptor Tim-3 in individuals with melanoma and non-muscle-invasive BCa ^14,19^. To characterize BCa-specific perturbations in NK cells more comprehensively, particularly across peripheral blood and tumor, we analyzed tumor tissue and peripheral blood mononuclear cells (PBMC) from a cohort of 68 BCa patients with either NMI and MI disease. We found that, in contrast to circulating blood NK cells, NK cells in the tumor microenvironment (TME) were functionally suppressed, and that TGF-β, and a hypoxia-dependent reduction in signaling through SLAM6 were responsible for this dysfunction. These results identify novel pathways of NK cell modulation that may have an impact on BCa progression, and that may be targetable to improve NK cell surveillance.

## Results

### Bladder cancer induces NK cell maturation in peripheral blood but not tumor

The frequency and composition of tumor-infiltrating leukocytes (TIL) in solid tumors differs across malignancies and often across patients with the same disease and clinical stage ^20–22^. However, the frequency of tumor-infiltrating T cells, NK cells, and dendritic cells (DC) is correlated with improved response to immune checkpoint blockade and overall survival ^23–26^. Tumors at distal sites can also influence the phenotype of immune cells in peripheral blood ^19,27,28^. To determine if BCa alters the frequency of major immune lineages, we performed FACS analysis on the PBMC (NMI: n=30, MI: n=16, HD: n=17), and primary tumors (NMI: n=13, MI: n=22) of a cohort of patients that included 22 individuals from whom matched PBMC and tumor tissue was available (**Table S1**), (Gating strategy depicted in **Figure S1A-S1B).** There was no difference in the number of total number of PBMC/mL of blood in patients compared to healthy donors (HD) **(Figure 1A)**, nor in the total number of cells/mg tumor tissue across disease stage **(Figure 1B)**. However, the frequency of TIL, (live, CD45^+^, singlets) was significantly lower in MI versus NMI tumors (**Figure 1C).**

**Figure 1:**
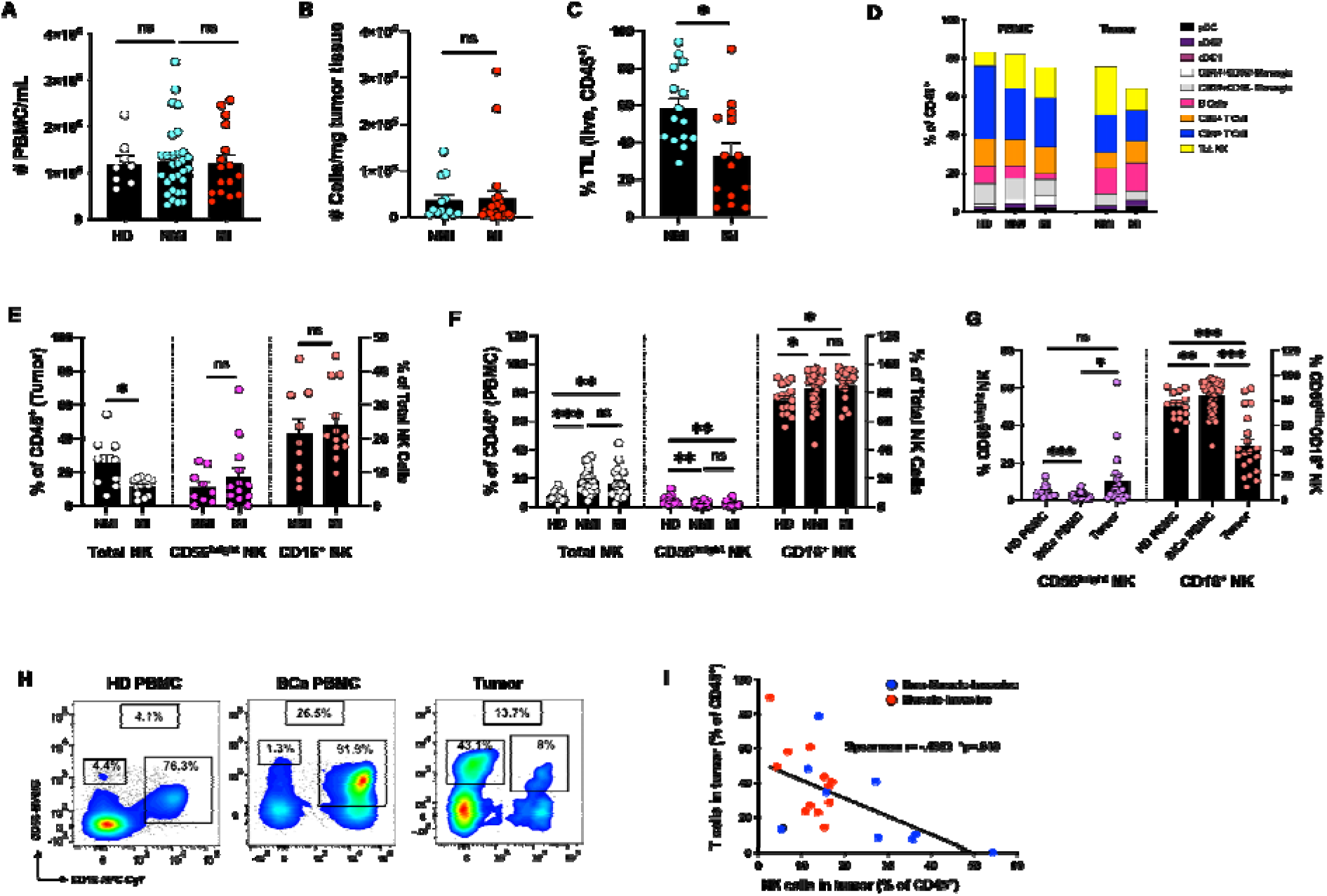
Bladder cancer induces NK cell maturation in peripheral blood but sustains NK cell immaturity in tumors. PBMC and cell suspensions obtained from dissociated tumor tissue were analyzed by flow cytometry to identify major immune lineages in BCa patients and HD. **(A)** Number of PBMC/mL of whole blood (HD: n=8, NMI: n= 30, MIs n= 22), **(B)** Number of viable cells/ mg of tumor tissue (NMI: n= 13, M: n= 22). **(C)** Frequency of TIL (live, CD45^+^ cells) in NMI and MI tumors NMI: n= 15, MI: n= 14). (**D)** Mean frequencies of major immune cell types in PBMC and TIL. **(E)** Frequency of NK cell populations in TIL expressed as a percentage of live, CD45^+^ singlets (NMI: n= 9, M : n= 13). Frequencies of CD56^bright^ and CD16^+^ sub-populations are plotted as as proportion of total NK cells on right axis. **(F)** Frequency of NK cell populations in the PBMC of HD and BCa patients (HD: n=16, NMI: n= 28, MI: n= 18). Frequencies of CD56^bright^ and CD16^+^ sub-populations are plotted as as proportion of total NK cells on right axis. **(G)** Comparison of the frequencies of CD56^bright^CD16^-^ and CD56^dim^CD16^+^ NK cells in PBMC and tumor. **(H)** Representative FACS plots of NK cells across tissue. Boxed numbers show the frequency of NK cells amongst CD45^+^ cells, while the percentages shown for each NK cell subset indicate the frequency amongst total NK cells. **(I)** Correlation between the frequencies of total NK cells and CD3^+^ T cells in bladder tumors (n=22). Significance for A-G was determined by unpaired Mann-Whitney test, with ns≥.05, *p<.05, **p <.01, ***p <.001. Straight line fitting for **I** was performed using least squares regression, and significance was determined by two-tailed Spearman correlation. Error bars show S.E.M. HD=Healthy Donor, NMI=Non-Muscle-Invasive, MI=Muscle-Invasive

NK cells were the second-most prevalent lineage identified in bladder tumors after T cells, comprising up to 25% of TIL in NMI tumors, but with significantly decreased infiltration of more advanced MI tumors where they comprised ∼11% of TIL (**Figures 1D** + 1E). Given the reported paucity of NK cells in normal bladder tissue^29,30^ and additional work showing infiltration of NK cells into bladder tumors^31^, these data suggest that NK cells recognize and infiltrate bladder tumors. Tumors, independent of stage, contained equivalent frequencies of monocytes (Figure S1C), dendritic cells (DC) (Figure S1D), B cells (Figure S1E), and CD4^+^ and CD8^+^ T cells (Figure S1F).

In peripheral blood there was a striking increase in the frequency of NK cells (pbNK) in BCa patients compared to HD, with NMI and MI patients having 2.4-fold and 2.2-fold increases respectively (**Figure 1F**). This accumulation was driven by an increase in more mature CD56^dim^CD16^+^ cells, and a concomitant decrease in less mature, CD56^bright^CD16^-^ NK cells. This suggests that increases in NK cell frequency and maturation in the periphery can be driven by a tumor at a distal site. BCa patient peripheral blood had no difference in the frequency of monocytes (Figure S1G) or DC (Figure S1H) compared to HD, but did have a significant reduction in the frequency of B cells in MI patients compared to HD (Figure S1I), and in CD4^+^ T cells independent of disease stage (Figure S1J).

Comparing NK cell subsets across tissue type revealed that tumor-infiltrating NK cells (tNK) are significantly less mature than pbNK from either HD or patients based on expression of the maturation marker CD16 (**Figure 1G**). CD16^+^ NK cells are considered as more cytolytic than CD16^-^ cells^32–34^ suggesting that tNK cells are not optimal for tumor surveillance. Representative FACS plots of pbNK from HD and BCa patients as well as tNK are shown in **Figure 1H**. Finally, there was a significant, inverse correlation, between the frequency of NK and T cells within the same tumor, with T cells being enriched in more advanced MI tumors (**Figure 1I**). This suggests that NK and T cells may respond differently to different TMEs associated with muscle invasion/non-invasion.

### Tumor-infiltrating NK cells are dysfunctional but not terminally so

We next determined if there were differences in effector function between NK cells from BCa patients and healthy donors via co-culture with K562 cells, a Class I HLA-deficient human leukemia cell line. pbNK cells from BCa patients produced IFNγ and degranulated, as determined by expression of LAMP-1, equivalently to pbNK cells from HD **(Figures 2A-2E)**. In contrast, tNK cells produced significantly less IFNγ compared to pbNK cells from both HD and BCa patients, and they were variably degranulated even in the absence of K562 targets **(Figures 2A-2E)**. Importantly, K562 targets were unable to elicit further degranulation **(Figures 2D + 2E**). Additionally, tNK cells released significantly less Granzymes A and B into culture supernatant after co-culture with K562 targets than pbNK cells **(Figure 2F)**. These results demonstrate that tNK cells are highly dysfunctional, likely due to repetitive degranulation within the tumor and additional TME-specific factors.

**Figure 2:**
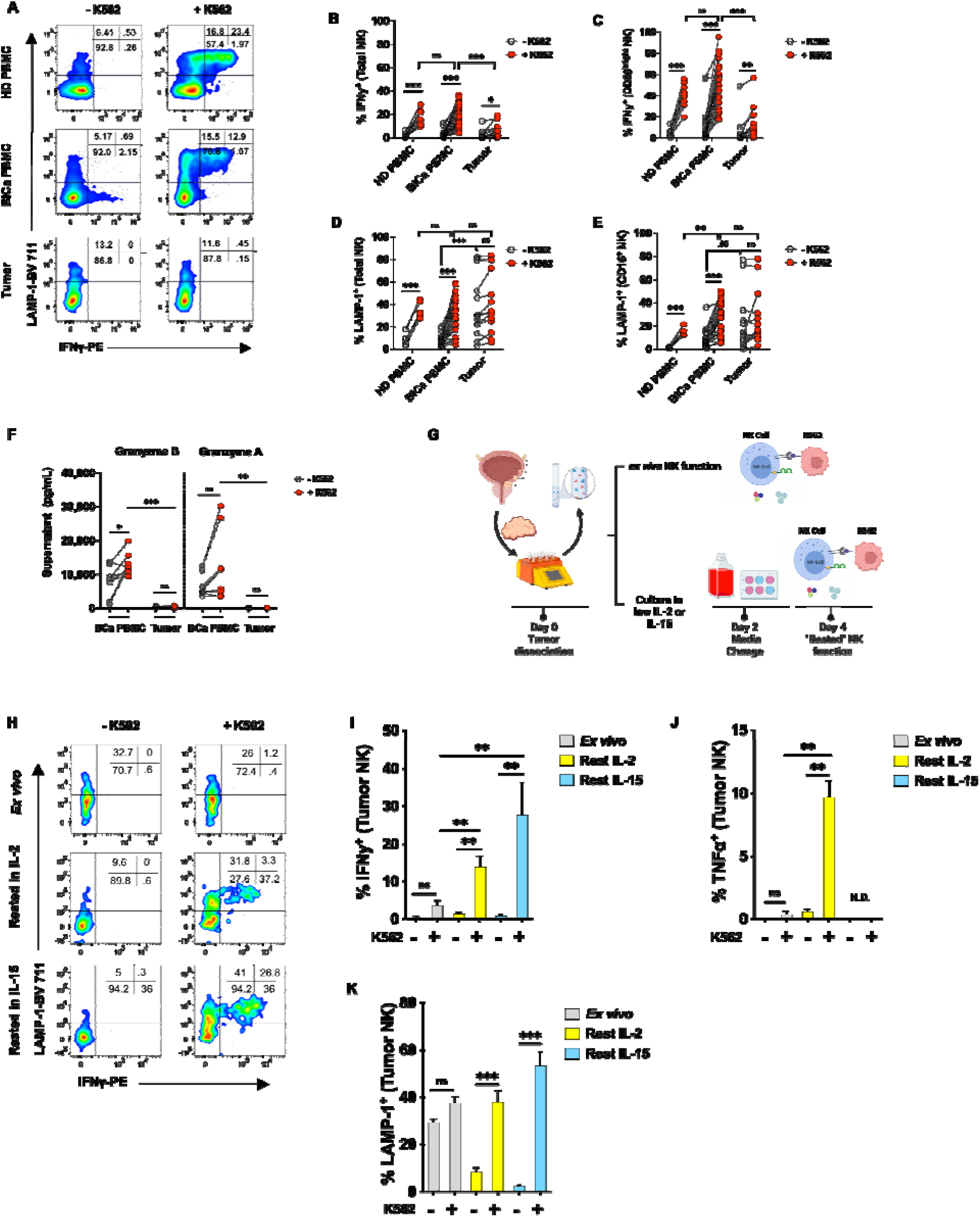
NK cells from the peripheral blood of BlCa patients are functionally competent while tumor-infiltrating NK cells are dysfunctional, but not terminally so. PBMC or single-cell suspensions obtained from primary tumors were activated overnight with 5ng/mL of IL-15 and co-cultured ± K562 target cells to induce IFNγ and degranulation. **(A)** Representative FACS plots of LAMP-1 and IFNγ gated on total NK cells from HD and BlCa patient PBMC, and from bladder tumors. **(B**) IFNγ production by total NK cells and **(C)** CD56^bright^ NK cells, (HD PBMC: n=10, BlCa PBMC: n=30, tumor: n=16). **(D)** LAMP-1 expression by total NK cells and **(E)** CD16^+^ NK cells, (HD PBMC: n=7, BlCa PBMC: n=25, tumor: n=14).). **(F)** Granzymes B and A released into the supernatant by NK cells from BCa PBMC (n=8) or tumor (n=6) after c—culture with K562 target cells **(G)** Experimental setup for “resting’ tNK cells by culturing dissociated tumor in 50U/mL IL-2 (n=10) or 1ng/mL IL-15 (n=3) for 4 days prior to K562 co-culture. **(H)** Representative FACS plots of IFNγ and LAMP-1 gated on total tNK cells after co-culture with K562 cells directly *ex vivo* (top), or after 4d culture in the presence of IL-2 (middle) or IL-15 (bottom). Frequencies of **(I)** IFNγ^+^, **(J)** TNFα^+^, or **(K)** LAMP-1^+^ tNK cells *ex vivo* (NMI=1, MI=2), or after 4 day culture in IL-2 (NMI=5, MI=5) or IL-15 (NMI=2, MI=1). All co-cultures were stained for IFNγ and LAMP-1, but only cells rested in IL-2 were stained for TNFα. . Error bars show S.E.M. Significance was determined by t-test, with ns≥.05, *p<.05, **p <.01, ***p <.001.

The dichotomy in effector function between pbNK and tNK cells suggested that suppressive elements unique to the TME undermine NK cell tumor surveillance. The architecture of the TME supports diverse suppressive strategies that can be mediated by soluble factors ^35–38^, cell-cell interactions^39,40^, and the creation of a metabolically hostile milieu^41,42^ To determine if physical disruption of the bladder TME ameliorated NK cell function, we dissociated tumor tissue, divided samples in half, and assessed NK cell function immediately *ex vivo*, or after 4 days of culture in media containing low doses of IL-2 (50U/mL) or IL-15 (1ng/nL) to ensure NK cell survival **(Figure 2G)**. Unexpectedly, NK cells challenged with K562 targets after 4 days of this “rest” culture regained the ability to produce IFNγ and TNFl1, and to degranulate appropriately **(Figure 2H-2K)**. The variable LAMP-1 expression observed in the absence of target cells *ex vivo* was also reduced to baseline, with LAMP-1 only expressed after addition of target cells **(Figure 2H + 2Ks)**. These data suggest that tNK cells are not terminally dysfunctional, and that a structured TME is required to undermine NK cell function.

### Inhibitory receptors expressed by tumor-infiltrating NK cells are detectable on NK cells in the peripheral blood

Immune Checkpoint Blockade (ICB) directed against Tim-3, TIGIT, and NKG2A is being evaluated in clinical trials for various solid tumors (e.g. NCT04139902, NCT04995523, and NCT05162755), and anti-PD-1/PD-L1 mAbs are already approved for many cancers including BCa^2,3^. However, most studies have focused on the effect of these mAbs in the context of relieving T cell exhaustion, despite significant promiscuity in receptor expression by both T and NK cells ^19,43–46^. To determine if NK cells express Tim-3, TIGIT, and/or NKG2A in the context of BCa, and if expression explained NK cell dysfunction in the tumor, we analyzed PBMC (NMI: n=30, MI: n=16, HD: n=17) and primary tumors (NMI: n=13, MI: n=22) from an expanded cohort of BCa patients that included 24 individuals from whom matched blood and tumor tissue were available. In bladder tumors we observed a stage-specific increase in Tim-3 expression by NK cells from NMI to MI tumors, whereas TIGIT was expressed equivalently regardless of disease stage (range 17-48%, **Figure 3A).** There was minimal expression of PD-1 by tNK cells as expected^47^ (range 2.7-5.9%, **Figure 3A)**. Identical expression patterns for Tim-3 and TIGIT were also observed on tumor-infiltrating CD4^+^ and CD8^+^ T cells, though PD-1 expression was higher than what was observed for NK cells (CD4^+^ T cells: 18-25.2% PD-1^+^, CD8^+^ T cells: 15.9-24.9% PD-1^+^) **(Figure S2A)**.

**Figure 3:**
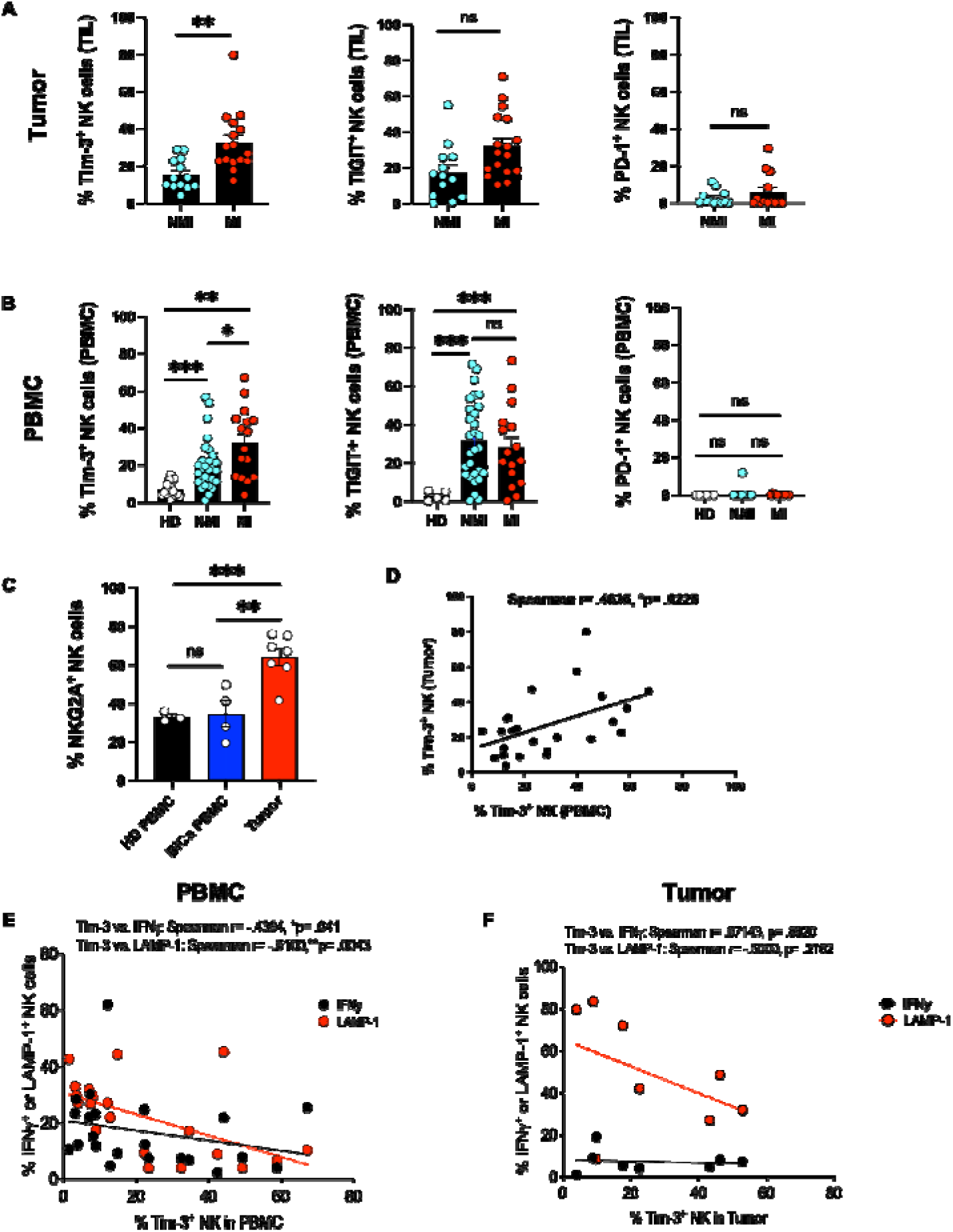
NK cells from BCa PBMC and tumors over-express inhibitory receptors. **(A)** Percentage of NK cells in tumor expressing Tim-3, TIGIT, or PD-1 (NMI: n=13, MI: n=16). **(B)** Percentage of NK cells in PBMC expressing Tim-3, TIGIT, or PD-1 (HD; n=17, NMI; n=29, MI; n=16). **(C)** Frequency of NKG2A expression on NK cells from HD PBMC (n=3) BCa PBMC (n=4) and TIL (n=7). **(D)** Correlation between the frequency of Tim-3^+^ NK cells in the matched PBMC and tumors of 24 BCa patients. **(E)** Scatter plot showing inverse correlation between Tim-3 expression on pbNK cells and IFNγ production or degranulation (n=22). **(F)** Scatter plot showing lack of correlation between Tim-3 expression on tNK cells and IFNγ production or degranulation (n=8). Significance for A-C was determined by unpaired Mann-Whitney test, with ns≥.05, *p<.05, **p <.01, ***p <.001. Straight line fitting for **D** was performed using least squares regression, and significance was determined by two-tailed Spearman correlation. Significance for **E** + **F** was determined by two-ailed Spearman correlation. Curve-fitting (dashed line) was performed using non-linear regression and a sum of squares F test. Error bars show S.E.M. HD=Healthy Donor, NMI=Non-Muscle-Invasive, MI=Muscle-Invasive.

In peripheral blood there was also a stage-specific increase in Tim-3 expression between NMI and MI patients similar to that observed for tNK cells (1.6-fold), as well as over-expression compared to HD pbNK cells (2.8-4.6-fold) (**Figure 3B).** TIGIT was also over-expressed by BCa pbNK cells relative to HD NK cells (10.8-11.8-fold), though again as found for tNK cells, in a stage-independent manner **(Figure 3B).** There was negligible PD-1 expression by pbNK cells **(Figure 3B)**. We observed similar expression patterns for Tim-3 TIGIT and PD-1 by pbCD4^+^ and pbCD8^+^ T cells **(Figure S2B)**. NKG2A is another inhibitory receptor recognizing HLA-E on target cells.^36,48^.. Examination of NKG2A in a subset of our cohort (HD PBMC: n=3, BCa PBMC: n=4, Tumor: n=7) revealed that tNK cells significantly over-express NKG2A relative to pbNK cells from HD and BCa patients **(Figure 3C and S2D) (Figure)**. tCD8^+^ T cells also over-expressed NKG2A relative to those in peripheral blood **(Figure S2C + S2D)**.

The unique disease stage-specificity of Tim-3 expression by NK cells across tissue suggested that pbNK cells may be useful as a readout of tumor invasiveness. Indeed, we found Tim-3 expression on NK cells from matched blood and tumor to be directly correlated **(Figure 3D),** maintaining this possibility. However, this correlation did not explain the dramatic difference in functional capacity between NK cells from the periphery and tumor. . We therefore compared Tim-3 expression with the ability of pbNK and tNK cells to produce IFNγ and degranulate. Increasing Tim-3 positivity was inversely correlated with the ability to degranulate and produce IFNγ by pbNK cells **(Figure 3E),** but not by tNK cells **(Figure 3F).** This suggests that over-expression of Tim-3 may be sufficient to suppress NK cell function, but that in the bladder TME it is not necessary.

### Ex vivo blockade of Tim-3 enhances IFNy production and release of granzymes from NK cells in BCa patient peripheral blood but not tumor

Pre-clinical blockade of Tim-3 and TIGIT with mAbs has been reported as enhancing NK and T cell function in mice and primary human cells ^43,46,49–52^. We tested if this strategy would improve the function of BCa pbNK and tNK cells by treating PBMC or dissociated tumor with commercially available anti-Tim-3 and anti-TIGIT mAbs (Tim-3: F38-2E2, TIGIT: MBSA34, and 741170) for 1h prior to, and during, K562 co-culture. Tim-3 blockade increased the frequency of total and CD56^bright^ IFNγ^+^ pbNK cells compared to control IgG **(Figures S3A-S3C)**, had no effect on LAMP-1/degranulation **(Figure S3A + S3D)**, but did enhance the release of Granzymes A and B **(Figure S3E)** .TIGIT blockade did not enhance any functional readout, and blockade of both receptors simultaneously resembled blockade of Tim-3 alone **(Figure S3F-S3H).** Importantly, blockade of neither receptor improved the function of NK cells from tumor, suggesting that dysfunctional NK cells in the TME may be marked by expression of inhibitory receptors, but their dysfunction is not caused by it.

### scRNASeq identifies distinct transcriptional differences between NK cells in peripheral blood versus bladder tumors

Having established that inhibitory receptor expression alone did not explain the severe dysfunction of tumor-infiltrating NK cells we performed 3’ single-cell RNA sequencing (scRNASeq) on live, CD45^+^ cells sorted from the PBMC (n=7) and tumors (n=6) of BCa patients to identify the TME-specific factors that suppress NK function **(Figure 4A and Table S3)**. As a comparison, HD NK cells were negatively bead-isolated from HD PBMC (n=5) and also sequenced. Details pertaining to scRNASeq QC are shown in **Figure S4A-S4C**. Single-cell libraries from all 18 samples, comprising 68,192 immune cells, were integrated using Seurat ^53,54^ to identify cell types common across tissue of origin and disease state. Graph-based clustering resulted in 19 clusters at resolution 0.8 **(Figure S4D)**, with 18,762 cells from four clusters (3, 4, 10, and 13) annotated as NK cells using module scores derived from published NK cell gene signatures^26,55–58^ **(Figures S4E + S4F and Table S4)**. The NK cells were sub-clustered against each other, resulting in 5 clusters of NK cells at resolution 0.2 for analysis **(Figure 4B)**. These NK cells were derived from each of the three tissues examined (HD PBMC: 7,809 NK cells, BCa PBMC: 7,082 NK cells, and TIL: 3,871 NK cells **(Figure S4G)** and represented each of the twelve individual donors **(Figure S4H) (Figure 4B)**. We then calculated the differentially expressed genes (DEG) in the NK cells from each cluster, parsed by tissue of origin **(Figure 4C)**. There was little overlap in gene expression by NK cells belonging to the same cluster but originating in different tissues. For example, in all clusters, genes defining NK cell effector signaling (CD247), and glycolysis (LDHA, ENO1), were significantly higher in HD pbNK cells compared to those from BCa patients regardless of tissue. Conversely, gene expression by tNK cells was associated with sustained activation (HLA-DRA), tissue-residency ^59^ (CD69^60^ and RGS1^61^), cellular stress (heat shock proteins, TSC22D3^62^), inhibition of signaling (NFKBI, DUSP1), and production of inflammatory cytokines (CCL3, CCL4, IFNG), but with little expression of cytolytic genes. Finally, pbNK cells from BCa patients were enriched in genes involved in lymphocyte trafficking (S1PR4, SELL), protein synthesis (ribosomal genes), and effector function (GZMH), but also inhibitory receptors (KLRG1). Importantly, this analysis allowed for identification of genes that may be specific to tNK cells, such as AREG, HLA-DRA, RGS1^61^ and CD69 **(Figure 4C)**, and revealed that both the presence of BCa, and the tissue of origin, induce specific programs of NK cell gene expression.

**Figure 4:**
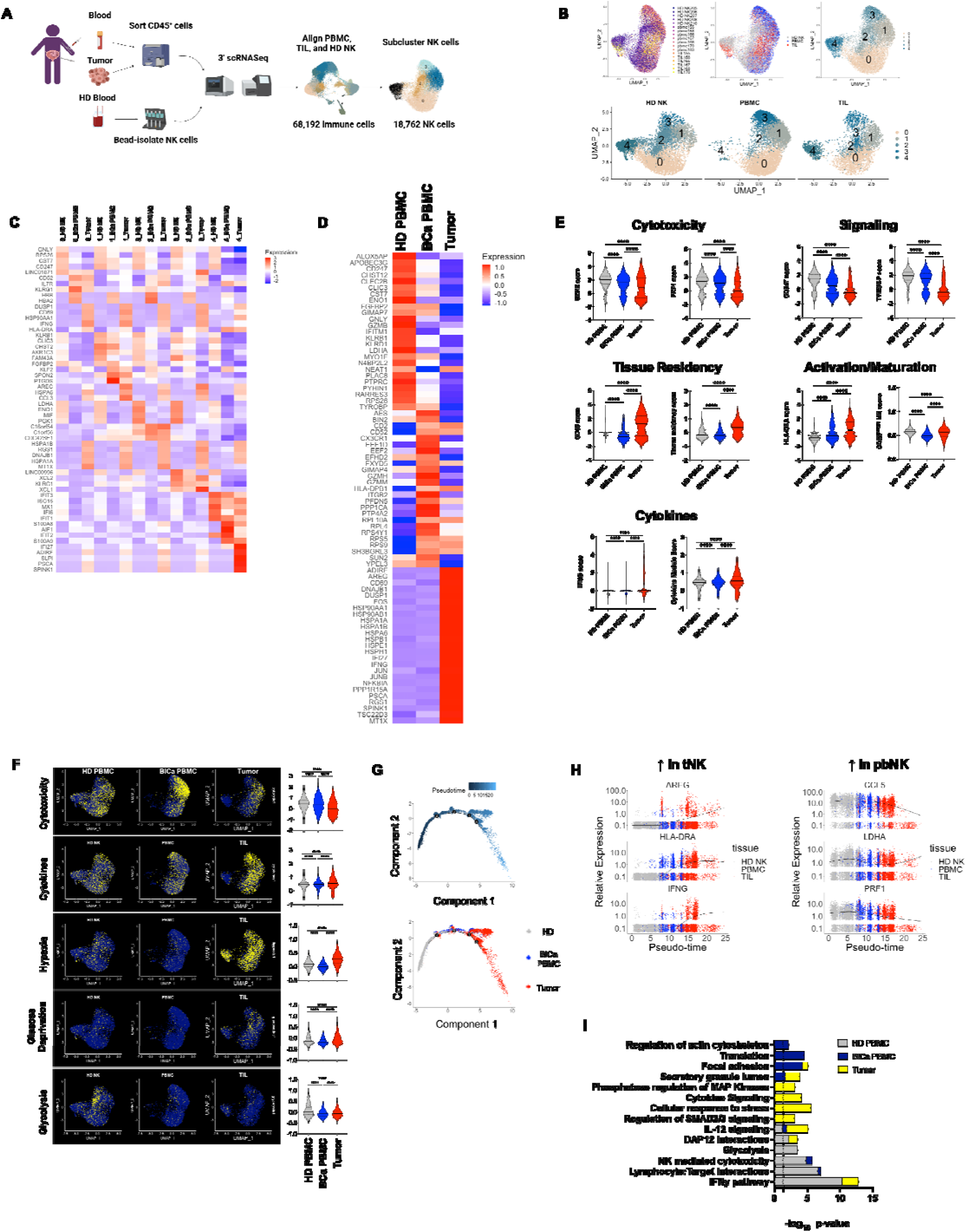
Distinct programs of gene expression define NK cells across tissue and disease state. **(A)** Total PBMC (NMI; n=2, MI; n=5) and patient-matched CD45^+^ TIL (NMI; n=2, MI; n=4) were FACS sorted and sequenced for 3’ scRNASeq usings the 10x platform. The resulting clusters were integrated with scRNASeq libraries generated from NK cells in HD PBMC (n=5) to allow comparisons across tissue and disease-state. **(B)** UMAP depicting 18,762 sub-clustered NK cells derived from bladder tumor, PBMC from BlCa patients, or HD PBMC. Clusters are colored by donor, tissue of origin, and cluster (top panel), and cluster membership separated by tissue (bottom panel). **(C)** Heatmap showing scaled expression of the top 5 significant DEG by cluster and tissue of origin, ordered by adjusted p-value, in pbNK cells from HD and pbNK and tNK cells from BCa patients. **(D)** Heatmap showing scaled expression of the top 25 significant DEG by tissue of origin, ordered by adjusted p-value, in pbNK cells from HD and pbNK and tNK cells from BCa patients. **(E)** Scaled expression of the indicated genes or gene signatures in NK cells from the indicated tissues. **(F)** Single-cell module scores for gene signatures pertaining to metabolic and immune processes across tissue of origin overlaid on UMAP plots (left) and quantified (right). For UMAP plots, non-zero expression between the 10^th^ and 90^th^ quantiles is shown. **(G)** Single-cell trajectory of NK cells colored by pseudotime (top) and by tissue of origin (bottom). **(H)** Expression of select genes as a function of pseudotime. **(I)** Select Gene Ontology and KEGG terms enriched in NK cells from peripheral blood or tumor. For violin plots, median expression (solid line) and quartiles (dotted lines) are shown with significance calculated using an unpaired Mann-Whitney t-test (p≥.05, *p<.05, **p <.01, ***p <.001, ****p <.0001). Significant differential gene expression was performed in Seurat using a non-parameteric Wilcoxon rank sum test. Pathway enrichment analysis was performed with Enrichr using Fisher’s exact test.

To examine the influence of both BCa and tissue type on NK cell gene expression, we compared significant DEG for NK cells according to tissue, **(Figure 4D)**, as well as by genes of interest **(Figure 4E)**. HD pbNK cells were significantly enriched in cytolytic (GNLY, CST7, FGFBP2, GZMB), and glycolytic (LDHA, ENO1) transcripts compared to NK cells from BCa patients, regardless of tissue. Patient pbNK cells expressed genes associated with cytoskeletal rearrangements and target cell adhesion (SH3BGRL3^63^, CD2^64^), Granzyme B-independent cytolytic activity (GZMH, GZMM), translation (ribosomal genes), and chemotaxis (S1PR1^65^, CX3CR1^66^) . In contrast, tNK cells expressed a paucity of cytolytic genes, were highly activated (CD69, HLA-DRA), immature (CD56^bright^ signature^67^), imprinted with tissue residency^61,68^ (CD69, CXCR6, ITGA1, ITGAE, RGS1), and had higher expression of a cytokine module that included IFNG (IFNG, TNF, CCL5, XCL1/2, FLT3LG) **(Figures 4C + 4E** and **Table S4**)

NK cell effector functions are directly linked to their ability to perform glycolysis ^69–71^. The dearth of glycolytic genes such as LDHA and ENO1, observed in patient versus HD NK cells (**Figures 4C + 4D**) prompted us to further examine metabolic perturbations in the bladder TME. Indeed, tNK cells had significantly higher scores for gene signatures associated with hypoxia ^72,73^ and glucose deprivation ^74^, as well as a lower glycolysis ^75^ score, compared to pbNK cells from both healthy donors and BCa patients **(Figure 4F and Table S4)**. Notably, these aberrations in tNK cells coincided with a lower cytotoxicity score ^76^, but higher expression of genes in a cytokine module comprised of IFNG, TNF, XCL1/2, and FLT3LG **(Figure 4F)**. These data suggested that the impaired cytolytic capacity of tNK cells might result from a sub-optimal metabolic milieu that is unique to the TME.

Given the general competence of pbNK cells we sought to determine if NK cells progress through a linear trajectory of increasing dysfunction, beginning with pbNK cells from HD, and terminating in tNK cells. Accordingly, we constructed single-cell pseudotime trajectories using Monocle^77^. This approach identified a continuum of NK cell differentiation with pbNK cells from HD at one terminus, tNK cells at the other, and pbNK cells from BCa patients in the middle **(Figure 4G)**. The expression of transcripts enriched in the tumor (e.g. AREG (epidermal growth factor-like protein amphiregulin), HLA-DRA, and IFNG) showed progressive up-regulation as pseudotime progressed, with low expression in healthy peripheral blood, induction in BCa patient peripheral blood, and maximal expression in tumor **(Figure 4H)**. Conversely, transcripts such as LDHA, the DC chemo-attractant CCL5, and the cytolytic gene perforin (PRF1), were maximally expressed in HD pbNK cells, declined in BCa pbNK cells, and reached lowest expression in sub-populations of tNK cells **(Figure 4H)**.

Finally, we conducted pathway enrichment analysis using Enrichr^78^, taking the top 100 significantly DEG from NK cells in each tissue as input. Effector pathways associated with NK cell surveillance (e.g. IFNγ, cytotoxicity, and lymphocyte-target cell interactions), as well as glycolysis were enriched in pbNK cells from HD, but largely absent in NK cells from BCa patients **(Figure 4I)**. pbNK cells from BCa patients were enriched in processes involving translation, focal adhesion, and regulation of the actin cytoskeleton, likely representing activated, migratory cells that are not yet dysfunctional. Gene expression by tNK cells was unique and involved processes associated with response to, and production of, cytokines such as IL-12 and IFNγ, but also SMAD2/3 signaling downstream of TGF-β, and phosphatase activity; two suppressive signatures that might underlie TME-specific NK cell dysfunction. Taken together the tissue-specific conservation of NK cell transcriptional programs mirrored the tissue-specific dysfunction we observed at the protein level. We chose to further examine TGF-β signaling, glucose deprivation, and hypoxia as mechanistically underlying tNK cell dysfunction because of their significance and specificity to the bladder TME.

### TGF-β suppresses NK cell effector function in bladder tumors

TGF-β is a pleiotropic cytokine that mediates suppressive effects on multiple hematopoietic cells in murine^79^ and human BCa^23^, including NK cells^59,80–84^. Because tNK cells were significantly enriched in the SMAD2/3 pathway that is a hallmark of signaling through TGF-β **(Figure 4I)** we examined expression of a gene set specifically comprising a TGF-β signaling signature, that was found to be associated with poor response to PD-L1 blockade in BCa^23^. Indeed, tNK cells were significantly enriched in gene expression downstream of TGF-β compared to pbNK cells regardless of BCa status **(Figure 5A).** To confirm this we tested the ability of TGF-β to inhibit effector function in HD NK cells. Culture of HD NK cells with 1ng/mL of IL-15 and TGF-β (.05ng/mL-50ng/mL) for 6 days significantly reduced both IFNγ and LAMP-1 expression after co-culture with K562 targets **(Figures 5B-5C)**. To increase phenotypic granularity we used the same culture system but performed CyTOF with an NK cell-focused panel (Table S2) on HD PBMC (n=20). After acquisition, PBMC were clustered with DepecheR^85^, and 4 NK cell clusters were annotated. Clusters 1 and 2 showed a significant reduction in IFNγ production and degranulation after exposure to TGF-β **(Figure 5D)**. TGF-β also inhibited IFNγ production and degranulation elicited by Antibody-Dependent Cellular Toxicity (ADCC) by using Raji cells coated with anti-CD20 as target cells instead of K562 **(Figure 5E)**. Cluster 1 NK cells were the most susceptible to inhibition for both assays and both effector readouts. In addition to being the largest Cluster, Cluster 1 NK cells expressed the most IdU (marking proliferative cells), CD56, Tim-3, and CD94 (co-receptor for NKG2-family receptors) **(Figure 5F)**. Cluster 2 NK cells were the second-most susceptible to TGF-β inhibition, and expressed the most Siglec-7, TIGIT and LILRB1, all inhibitory receptors, as well as CD16. This suggests that TGF-β may inhibit effector functions both in actively proliferating, less mature NK cells, as well as in more mature NK cells that have acquired an inhibitory phenotype.

**Figure 5:**
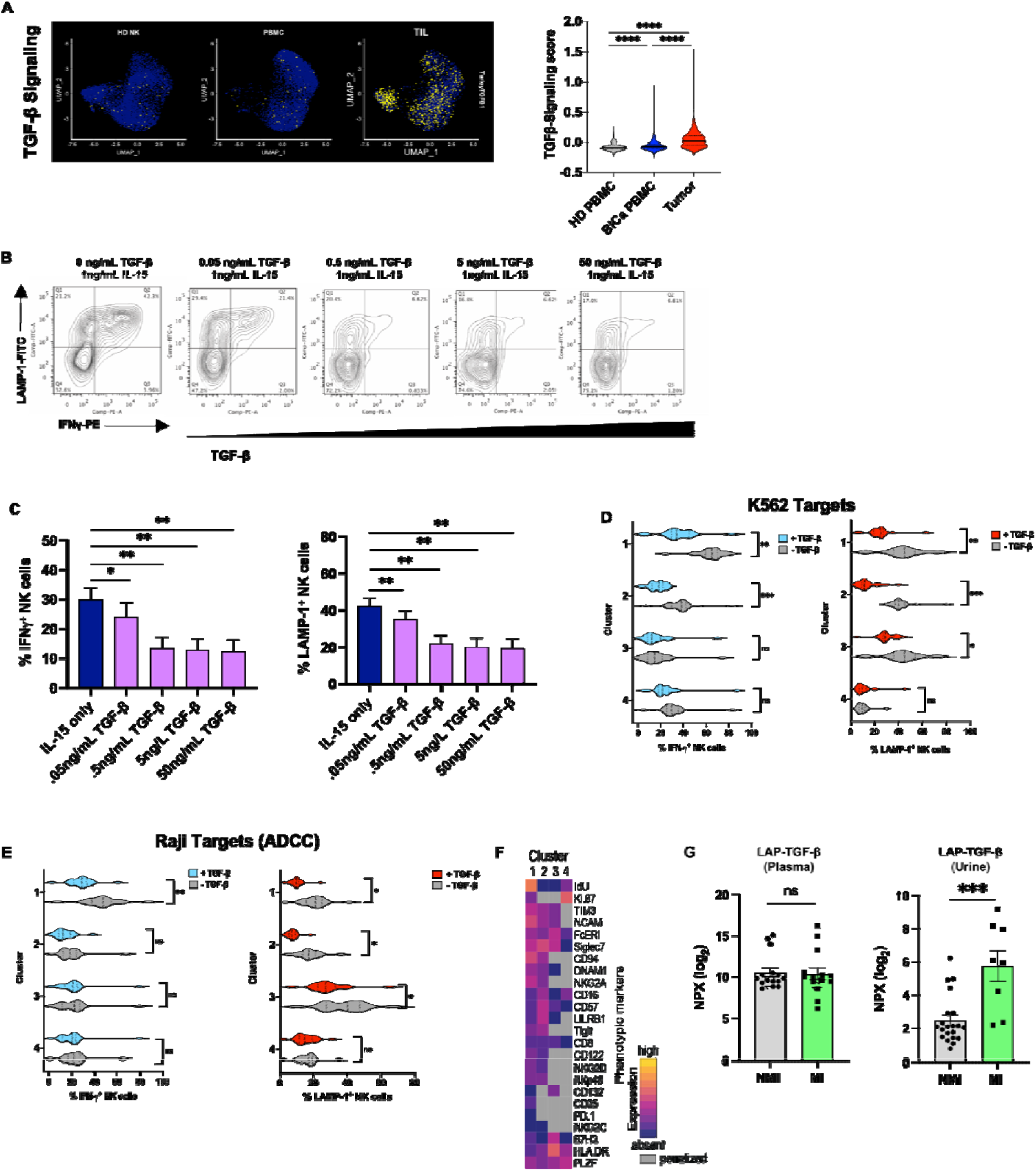
TGF-β in the TME suppresses NK cell function. **(A)** Single-cell module score for a TGF-β signaling signature overlaid on UMAP plots of pbNK cells from HD and BCa patients, or tNK cells (left) and quantified (right). For UMAP plots, non-zero expression between the 10^th^ and 90^th^ quantiles is shown. HD NK cells (n=6) were bead-isolated from PBMC and cultured for 6 days in the presence of 1ng/mL IL-15 alone or in combination with increasing concentrations of TGF-β (.05ng/mL-50ng/mL), before K562 target cell co-culture and FACS. **(B)** Representative FACS plots for IFNγ and LAMP-1. **(C)** Frequency of IFNγ^+^and LAMP-1^+^ NK cells from day 6 for IL-15 ± TGF-β-treated NK cells. **(D)** HD PBMC (n=20 donors) were cultured with 1ng/mL IL-15 or IL-15 and 5ng/mL TGF-TGF-β, for 6 days, analyzed by CyTOF, and clustered with Depeche. Comparison of IFNγ production and LAMP-1 expression, with or without TGF-β, for each cluster was performed. **(E)** HD NK cells (n=20) were cultured as above, but anti-CD20 coated Raji cells were used as targets to assess function elicited by ADCC. Comparison of IFNγ production and LAMP-1 expression, with or without TGF-β, for each cluster was performed. **(F)** Protein-level expression of markers for each cluster as determine by CyTOF. **(G)** Olink analysis of soluble proteins was performed on BCa patient plasma (NMI=15, MI=14) and cell-free patient urine (NMI=21, MI=8). Significance was calculated using paired t-test for **C** and unpaired Mann-Whitney t-test for **G.** Error bars show S.E.M .Significance for CyTOF was performed by . . . . . (p≥.05, *p<.05, **p <.01, ***p <.001, ****p <.0001).

To validate whether TGF-β was explicitly detectable in BCa patients we performed O-link analysis to profile soluble proteins in patient plasma and cell-free urine. There was no difference in circulating TGF-β in BCa plasma across disease stage **(Figure 5G)**, but there was significantly more TGF-β in the urine of MI patients compared to NMI **(Figure 5G).** Urine cytology is used clinically to identify tumor cells during diagnosis and follow-up, and has recently been shown to contain immune cells that better resemble those found in bladder tumors that those from peripheral blood, presumably due to contact between urine and the tumor^86^ (Tran and Bhardwaj *in revision).* While speculative, TGF-β in the TME likely acts to suppress NK cell function^81,87^, may reduce NK cell efficacy via reprogramming into less cytolytic ILC1^88^, and may contribute to the physical exclusion of NK cells from tumor nests as shown to occur for T cells in bladder tumors^23^.

### Hypoxia suppresses NK cell effector function in the bladder TME

The most striking differences in gene expression between pbNK cells and tNK were TME-specific metabolic disruptions such as defective glycolysis, hypoxia, and glucose deprivation **(Figure 4F)**. We sought to validate the latter two, using HD pbNK cells and the human IL-2-dependent NK cell line NK-92. First, we used the fluorescent glucose analog 2-NBDG to assess the ability of pbNK and tNK cells to uptake glucose. tNK cells had the ability to take up significantly more glucose than pbNK cells from both BCa patients and healthy donors **(Figure 6A)**. This suggests that the gene expression associated with glucose deprivation may reflect low levels of extracellular glucose in the TME rather than a defect in the ability to access it.

**Figure 6:**
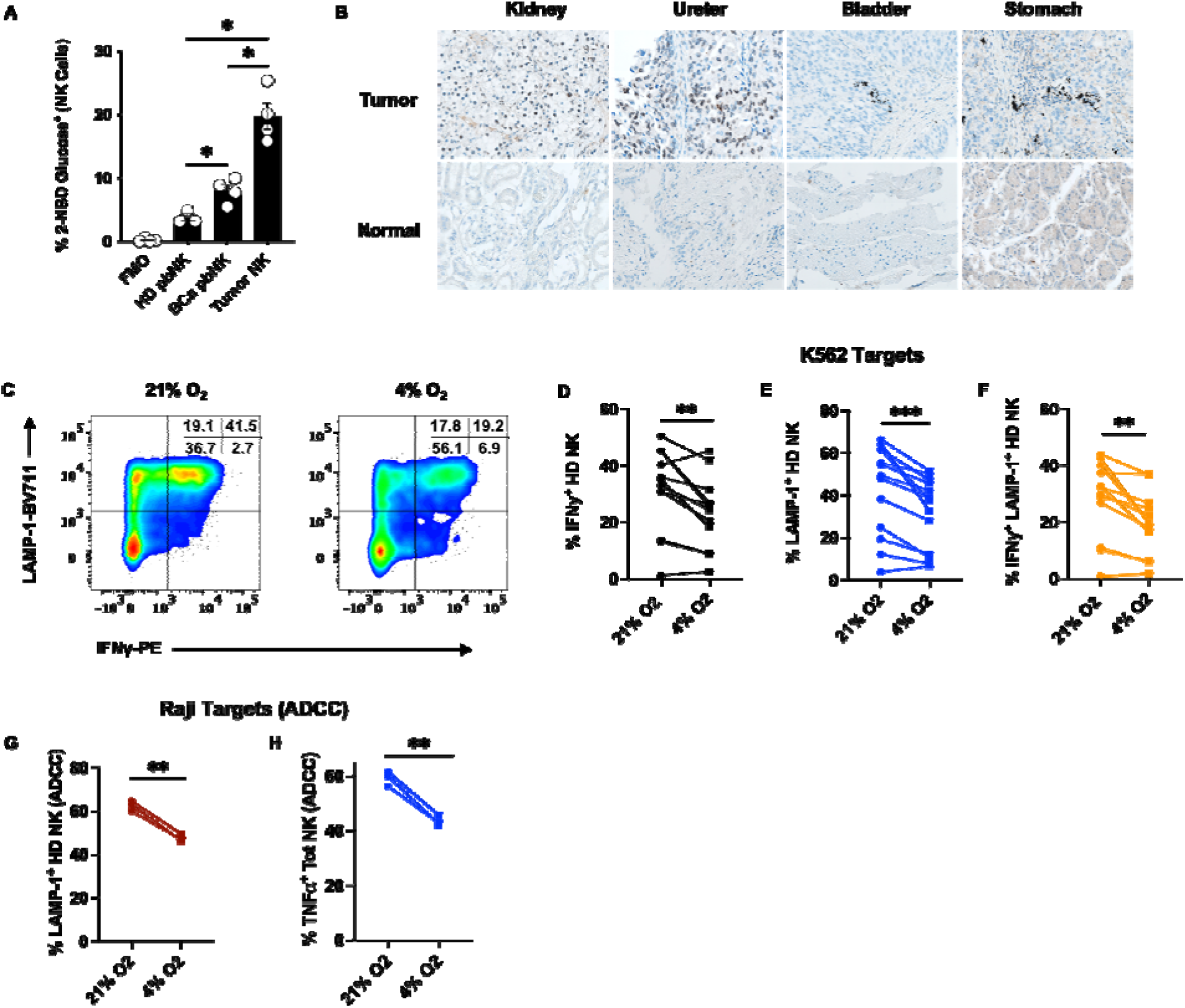
Hypoxia suppresses NK cell function. PBMC from HD (n=3), BCa patients (n=4) or dissociated tumor (n=4) were incubated with the fluorescent glucose analog 2-NBDG-FITC to determine glucose uptake, then stained with FACS antibodies to identify NK cells by flow cytometry. **(A)** Percentage of 2-NBDG^+^ NK cells. **(B)** FFPE sections of kidney, ureter, bladder, and stoma h tumors, as well as normal tissue from those organs (unmatched), was stained for HIF-111 (brown) and hematoxylin (blue). Images are magnified 63x and representative of two independent experiments. **(C)** Representative FACS plots of IFNγ and LAMP-1 on HD NK cells cultured for 3 days under normoxic (21% O_2_) or hypoxic (4% O_2_) conditions prior to co-culture with K562 targets. **(D)** Frequencies of IFNγ^+^, **(E)** LAMP-1^+^, or **(F)** IFNy^+^LAMP-1^+^ HD NK cells (n=13) after 3-4 days of normoxic or hypoxic culture followed by co-culture with K562 target cells. **(G)** ADCC-induced expression of LAMP-1 and **(H)** TNFα by HD NK cells (n=3) in response to anti-CD20 and co-culture with Raji cell targets after 4 days of normoxic or hypoxic culture. Significance was calculated using unpaired t-test for **A** and paired t-test for **D-H** with error bars showing S.E.M. and p≥.05, *p<.05, **p <.01, ***p <.001, ****p <.0001.

Gene expression associated with hypoxia was present almost exclusively in tNK cells **(Figure 4F)**. We performed IHC for HIF-1l1 on an FFPE tissue array containing sections of kidney, ureter, bladder, and stomach tumors, as well as normal tissue from those organs. HIF-1l1^+^ cells were present in all tumor sections but not normal tissue **(Figure 6B)** confirming the scRNAseq data. Next, we isolated HD pbNK cells and cultured them at 21% or 4% O_2_ for 3-4 days prior to co-culture with K562 targets and assessed their function. Hypoxic HD NK cells had significant reductions in IFNγ production and degranulation compared to those in normoxic culture **(Figures 6C-6F)**. Hypoxia also suppressed the ability of HD pbNK cells to degranulate **(Figure 6G)** and produce TNFα **(Figure 6H)** in response to Antibody Dependent Cellular Cytotoxicity (ADCC). Having confirmed that that low extracellular O_2_ in the TME inhibits NK cell surveillance, we next sought to determine the molecular mechanism by which this occurs.

### A hypoxia-induced defect in SLAMF6 signaling suppresses NK cell effector function in bladder tumors

To identify changes in NK cell signaling that occur during hypoxia we used a dot blot array to detect changes in the phosphorylation status of 59 relevant receptors containing ITAM/ITIM/ITSM (Immunoreceptor Tyrosine Activating/Inhibitory/Switch Motif). Bead-isolated HD NK cells were cultured at 21% or 4% O_2_ for 3-4 days and used to make protein lysates before incubation with capture Abs on the array, and subsequent probing with an HRP-conjugated anti-phospho-tyrosine Ab. Hypoxic culture reduced signaling through multiple activating receptors such as NKp46, NKp80, KIR2DL4, and DNAM-1 **(Figure 7A)**. However, the most striking difference was the reduction in phosphorylation of the ITSM of SLAMF6 (Signaling Lymphocyte Activation Molecule 6) after hypoxic culture **(Figures 7B + 7C).** SLAMF6 is a homotypic receptor (i.e., the ligand for SLAMF6 is also SLAMF6) expressed by T, B, and NK cells^89^. It functions as an activating receptor in human NK cells and mediates IFNγ production and degranulation via recruitment of the adaptor proteins SAP and EAT-2 to its phosphorylated ITSM.^89–93^ Importantly, when HD NK cells were cultured at 4% O_2_ for 3-4 days, then “rested” at 21% O_2_ for 3 days, phosphorylation of the ITSM of SLAMF6 was restored **(Figures 7B + 7C),** analogous to the restoration of effector function observed after “rest” of tNK cells from tumors. These data suggest that hypoxia suppresses both NK cell function and signaling through SLAMF6, and that restoration of normoxia is accompanied by restoration of SLAMF6 signaling and NK cell function. Indeed, gene expression of SLAMF6, SH2D1A (SAP), SH2D1B (EAT-2), PRF1 (Perforin), and GZMB (Granzyme B) were lowest in tNK cells, while tNK had the highest expression of PTPN6 (SHP-1), a phosphatase involved in inhibition of SLAMF6 signaling^89^ and ZNF683, a zinc finger protein shown to inhibit transcription of EAT-2 and suppress NK cell cytotoxicity^94^ **(Figure7D)**. Together, these results implicate a hypoxia-induced inhibition of SLAMF6 signaling as underlying tNK cell dysfunction.

**Figure 7:**
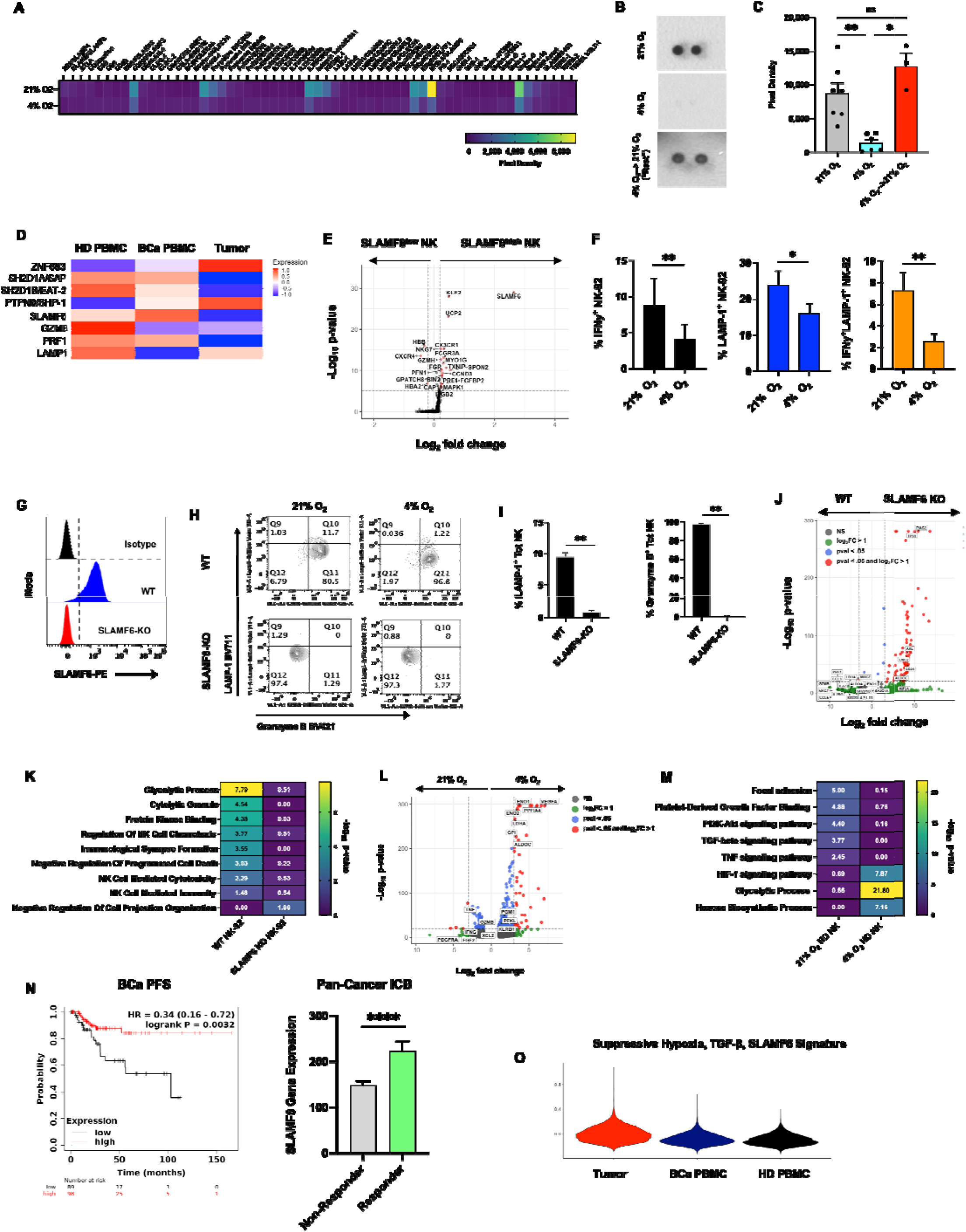
NK cell dysfunction in the TME results from a hypoxia-induced suppression of signaling through SLAMF6. **(A)** Densitometry quantification of tyrosine phosphorylation in the ITAM/ITIM/ITSM of 59 immune receptors on HD NK cells after 4 days of normoxic (n=5) or hypoxic culture (n=4). **(B)** Representative blot showing tyrosine phosphorylation on the ITSM of SLAMF6 in HD NK cells after 3-4 days of normoxic or hypoxic culture, or 3 days of hypoxic culture followed by 3 days of normoxic culture (“Rest”). **(C)** Densitometry quantification of tyrosine phosphorylation on the ITSM of SLAMF6 in HD NK cells after 3-4 days of normoxic (n=8) or hypoxic culture (n=6), or 3 days of hypoxic culture followed by 3 days of normoxic culture (“Rest”) (n=3). **(D)** Single-cell expression of genes involved in SLAMF6 signaling as well as genes for effector function in pbNK from HD and BCa patients, and tNK cells. **(E)** Volcano plot showing enrichment of effector gene expression in SLAMF6^high^ versus SLAMF6^low^ NK cells from scRNAseq (HD PBMC: n=5, BCa PBMC: n=7, Tumor: n=6). Expression of >1 was used as the threshold for SLAMF6^high^ NK cells. **(F)** Frequencies of for IFNγ^+^, LAMP-1^+^, or IFNy^+^LAMP-1^+^ NK-92 cells (n=3) after 3-4 days of normoxic or hypoxic culture and co-culture with K562 target cells. **(G)** Representative histograms showing SLAMF6 expression in WT NK-92 and RNP CRISPR-generated SLAMF6-KO NK-92 cells (n=5). **(H)** Representative FACS plots of LAMP-1 and Granzyme B expression in WT or SLAMF6-KO NK-92 cells after 3 days of normoxic or hypoxic culture followed by co-culture with K562 target cells. **(I)** Expression of Granzyme B and LAMP-1 by WT or SLAMF6-KO NK-92 cells after co-culture with K562 targets. Data are from two independent experiments. **(J)** Bulk RNAseq of WT or SLAMF6-KO NK-92 cells showing an enrichment of effector gene expression in WT cells compared to SLAMF6-KO. **(K)** Select Gene Ontology or KEGG terms significantly enriched in WT and SLAMF6 KO NK-92 cells. The top 100 significant DEG were used as input. **(L)** Bulk RNAseq of HD NK cells cultured at 21% or 4% O_2_ for 3 days. **(M)** Select Gene Ontology or KEGG terms significantly enriched in HD NK cells cultured at 21% or 4% O_2_ for 3 days. The top 100 significant DEG were used as input**. (N)** PFS for BCa patients stratified by high or low SLAMF6 gene expression as determined by bulkRNAseq (kmplot.com, n=187). **(O)** Expression of a gene signature comprised of inhibitory SLAMF6 pathway genes, hypoxia, and TGF-β signaling transcripts (Table S4) in scRNASeq of NK cells from the indicated tissues. Response to ICB stratified by SLAMF6 expression across 13 solid tumors in a cohort of 1,160 patients (kmplot.com). ITAM/ITIM/ITSM= Immunoreceptor Tyrosine Activating/Inhibitory/Switch Motif. PFS= Progression Free Survival, ICB= Immune Checkpoint Blockade Error bars show S.E.M. ns ≥.05 *p<.05, **p <.01, ***p <.001, ****p <.0001 as determined by unpaired or paired t-test

We next examined differences in gene expression between SLAMF6^low^ and SLAMF6^high^ NK cells in our scRNAseq data. This revealed that SLAMF6^low^ NK cells lacked effector transcripts involved in cytotoxicity (e.g. NKG7, GZMH, FCGR3A/CD16, and FGFBP2) compared to SLAMF6^high^ NK cells, and therefore that SLAMF6 is important for proper cytolytic function (**Figure 7E).** To test this directly, we used CRISPR to delete SLAMF6 in NK-92 cells, a human IL-2-dependent NK cell line, rather than primary NK cells which are technically more challenging to modify. We first validated that NK-92 cells were amenable to hypoxia-induced dysfunction analogously to primary NK cells and found that they were **(Figure 7F)**. Using a Ribo Nucleo Protein (RNP) approach and nucleofection efficiently deleted SLAMF6 at the protein level in NK-92 cells **(Figure 7G)**. Importantly, after co-culture of SLAMF6-KO cells with K562 targets there was significantly less LAMP-1 and Granzyme B expression compared to the mock control, confirming that an inability to signal through SLAMF6 compromises NK cell cytolytic function **(Figure 7H + 7I)**. As a control, co-culture with K562 cells at 4% O_2_ reduced degranulation by WT NK-92 cells as expected, and resulted in a further, but minimal reduction by SLAMF-KO cells **(Figure 7H)**. These results suggest that SLAMF6 is centrally involved in mediating NK cell cytolytic activity.

To identify any global changes in gene expression that accompany deletion of SLAMF6, we performed bulk RNAseq of WT or SLAMF6-KO NK-92 cells. SLAMF6-KO NK-92 cells showed significant downregulation of genes involved in cytolytic activity (PRF1, GZMB, NKG7), glycolysis (LDHA, ENO1, ALDOA1), and cytokines (CCL3, CCL5, IL32) **(Figure 7J)**), supporting our protein-level results from *ex vivo* tNK cells **(Figure 2A-2F)**. Additionally, as RNAseq was performed on normoxic NK-92 cells, it was notable that SLAMF6 positively regulated both cytolytic function and glycolysis. This was further supported by pathway analysis of the significant DEG between WT and KO NK-92 cells **(Figure 7K)**.

We also performed bulk RNAseq on HD NK cells cultured at 21% or 4% O_2_ for 3 days to directly query how hypoxia affects gene expression in primary NK cells. Hypoxic HD NK cells had significantly less expression of cytokine (TNF, IFNG, XLC2) and cytolytic (GZMB) transcripts than normoxic cells **(Figures 7L + 7M**), supporting our functional results. Conversely, hypoxic HD NK cells were enriched in genes involved in inhibitory signaling (KLRB1/CD161^95^, VEGFA^96^) HIF-1l1 signaling, and glycolysis **(Figures 7L + 7M)**, together describing a state of increased NK cell dysfunction in low oxygen, and again in harmony with our previous observations.

We next queried publicly available datasets to determine if SLAMF6 expression was associated with clinical outcomes^97,98^. Higher SLAMF6 expression was significantly correlated with longer progression-free survival in BCa patients (n=187), and improved response to ICB across 13 solid tumors including BCa, in a cohort of 1,160 individuals **(Figure 7N)**. Finally, a gene signature combining hypoxia and TGF-β and inhibitory SLAMF6 signaling was most enriched in tNK cells followed by BCa pbNK and HD pbNK cells **(Figure 7O and Table S4)**. This underscores the contribution of all three TME-specific axes in undermining NK cell surveillance in bladder tumors.

## Discussion

Our study provides the first comprehensive analysis of NK cell function in BCa patients, reveals important differences between pbNK and tNK cells, and identifies SLAMF6 as a new translationally relevant target for immunotherapy. We confirmed and expanded on previous work showing that bladder tumors are substantially infiltrated by NK cells, and that pbNK cells from BCa patients have no cytolytic impairment^31,99,100^. NK cells represented the second most abundant immune lineage after T cells in NMI tumors, suggesting that they detect early perturbations associated with malignancy and traffic to bladder tissue efficiently. We also observed a significant, inverse relationship between NK cell infiltration and tumor invasiveness. This suggests that NK cells are important for controlling tumor progression, but that increasing TGF-β, hypoxia, and defective signaling through SLAMF6 in the TME, ultimately subvert this as tumors progress.

Up-regulation of inhibitory receptors such as PD-1 is a hallmark of T cell dysfunction in cancer^18^. We identified a parallel process for NK cells that involved the inhibitory receptors Tim-3, TIGIT, and NKG2A, all of which have been correlated with disease progression and decreased overall survival in multiple cancers ^101,102^. Tim-3 was over-expressed both by pbNK and tNK cells in a disease stage-specific manner. This suggests that Tim-3^+^ pbNK cells have utility as a biomarker by which to assess disease progression. Conversely, TIGIT was over-expressed by pbNK and tNK cells independent of disease stage, making TIGIT^+^ NK cells useful as an indicator of the presence of BCa. These findings validate and extend our previous work showing up-regulation of Tim-3 and TIGIT by pbNK cells in NMI BCa patients^103^.

However, Tim-3 and TIGIT expression alone could not explain the TME-specific dysfunction of tNK cells. For example, while Tim-3 was expressed equivalently across both tissues, its expression was only inversely correlated with impaired NK function in peripheral blood, not tumor. Additionally, while *ex vivo* blockade of Tim-3 enhanced IFNy production and release of Granzymes A and B by pbNK cells, there was no synergy when combined with TIGIT blockade, and neither improved tNK cell function. Efforts to improve tNK cell function with a commercially available SLAMF6-Fc chimeric protein, were not successful (Data not shown). One interpretation of these results is that commercial clones simply do not block signaling through TIGIT. Indeed, multiple studies showing enhanced NK or T cell function after TIGIT blockade have been conducted using murine cells^45,59^ proprietary anti-TIGIT Abs^46,59^, combining TIGIT blockade with blockade of PD-(L)1^59,104,105^, and/or humanized anti-TIGIT Abs (REF). Alternatively, the lack of efficacy of both Abs for tNK cells may reflect a lack of ligand expression in the TME or an inability of Ab-based blockade to overcome suppression imprinted by the bladder TME, mediated by hypoxia, glucose deprivation, TGF-β-signaling, and likely other factors.

To determine what caused TME-specific NK cell dysfunction, barring inhibitory receptor expression, we performed scRNAseq of matched BCa PBMC and TIL, as well as HD pbNK cells. While tNK cells were significantly enriched in transcripts associated with activation (CD69, HLA-DRA) and cytokine production (IFNγ, TNFl1, XCL1/2, FLT3LG), they were simultaneously enriched in genes involved in TGF-β signaling (SMAD2/3), cellular stress, hypoxia, and glucose deprivation, as well as possessing a gene signature implicating deficient signaling through SLAMF6. Indeed, metabolic dysregulation and TGF-β in the TME have been shown to subvert cytolytic activity ^106,107^, and TGF-β specifically can additionally convert tNK cells into poorly cytolytic Type 1 innate lymphoid cells^88,108^, as well as directly suppress NK cell glycolysis and by extension, cytolytic function^109,110^. We validated the ability of TGF-β to suppress IFNγ production and degranulation by HD pbNK cells, both in response to K562 targets as well as when triggered by ADCC. Specifically, the largest NK cluster comprised of less mature, proliferative cells, was also the most sensitive to TGF-β. This implies that not only are most NK cells sensitive to TGF-β-mediated suppression in the TME, but that impairment of this population specifically would interfere with maturation towards a more cytolytic CD16^+^ population, and possibly reduce the number of NK cells in bladder tumors. These results, along with the increased amount of LAP-TGF-β present in urine from MI patients compared to NMI, support a strong, TME-specific, suppressive role for TGF-β in undermining tNK cell function in bladder cancer.

We also confirmed our scRNAseq data proposing hypoxia as a putative, TME-specific, inhibitor of NK cell function. Hypoxic culture significantly suppressed IFNγ and TNFα production, and decreased degranulation by competent pbNK cells in response to both K562 targets and elicited by ADCC. HIF-1α was also detectable by IHC in bladder tumor but not normal bladder. Furthermore, hypoxia, even in the absence of additional TME-specific suppressive axes like TGF-β and glucose deprivation, suppressed signaling through multiple NK cell activating receptors such as DNAM-1, NKp46, NKp80, and SLAMF6. While speculative, a global inability to translate activating signals from the TME into effector function would have deleterious consequences for NK cell surveillance in bladder tumors.

These results support literature showing improved function and tumor clearance in HIF-1α^-/-^ NK cells^111^. We also showed that deletion of SLAMF6 abrogated NK cytolytic activity and production of Granzyme B even under normoxic conditions, suggesting that SLAMF6 is critical to the ability of NK cells to kill. Combined with our results demonstrating significantly less signaling through SLAMF6 during hypoxic culture, this suggests that low extracellular O_2_ alone may cripple the ability of NK cells to destroy tumor cell targets in the TME. Finally, deletion of SLAMF6 also impaired glycolytic processes, again, even under normoxic conditions, suggesting that it may have a role in NK cell metabolism as well as NK cell cytotoxicity. Alternatively, it may be that the inability to signal through SLAMF6 reduces glycolytic capacity which in turn limits cytolytic activity.

Importantly however, we observed normoxic levels of phosphorylation in the ITSM of SLAMF6 after “resting” hypoxic NK cells for several days in normoxic conditions. As we also observed restoration of all tNK cell effector functions after “resting” bladder tumor in normoxic conditions, it is likely that functional recovery from hypoxia is mediated by restoration of SLAMF6 signaling, as well as an overall increase in aerobic respiration While speculative, some degree of NK cell proliferation in “rested” tumor cultures, in response to the low IL-2 /IL-15 added, could result in the presence of functional daughter cells by the end of the “rest” period. However, we expect that restored signaling through SLAMF6 plays a more significant role, as protocols aimed at inducing NK proliferation typically use much higher cytokine concentrations, and expand NK cells over the course of weeks, not days^112–114^.

As mentioned above, bulk RNAseq of SLAMF6-KO cells revealed that SLAMF6 was also required for normal glycolysis, which has not previously been reported to our knowledge. A recent study showed that stem-like, partially-exhausted LCMV-specific CD8 T cells that were HIF-1l1^-/-^, over-expressed SLAMF6^115^ and *Tcf7*, suggesting that the inability to “detect” hypoxia maintained SLAMF6 expression. This is consistent with our observation that SLAMF6 expression decreases in the hypoxic TME. NK cells have higher basal rates of glycolytic activity than other lymphocytes so as to immediately respond to target cells^69,110^. The dramatic lack of cytolytic function in SLAMF6 KO cells is partially due to its described role in mediating NK effector function^89,91,93,116^, but may also result from a role for SLAMF6 in modulating glycolysis.

Our results are consistent with work demonstrating that SLAMF6 functions as an activating receptor on human NK cells, with its deletion reducing NK killing of non-hematopoietic target cells^90^, its recognition of influenza hemagglutinin and subsequent induction of NK cell function^117^, and the ability of rSLAMF6 to enhance NK cell proliferation and IFNγ production^118^. SLAMF6 expression by CD8^+^ T cells also marks the proliferative, progenitor-exhausted, stem-like cells that contribute to response to PD-1 blockade^119,120^, but has also been described as having both co-activating^121–123^ and co-inhibitory function^122,124^ in a context-dependent manner.

The clinical relevance of competent SLAMF6 signaling is suggested by the improved progression-free survival in BCa patients, and the improved response to ICB in individuals with various solid tumors including BCa, in those with higher rather than lower expression. While this data does not implicate SLAFM6 expression specifically on NK cells as being protective, SLAMF6 is overwhelmingly expressed by NK, T and B cell populations, so it is likely that higher expression confers at least a general advantage in tumor immunosurveillance.

This work reveals dramatic, tissue-specific differences in the function, gene expression, signaling, and amenability to ICB, of NK cells in the context of BCa. Additionally, we identified a novel mechanism by which hypoxia specifically suppresses NK cell function in the TME by abrogating signaling through SLAMF6. Interventions aimed at ameliorating the metabolically-hostile TME and enhancing signaling through pathways suppressed by it, will hopefully have translational benefit for patients with multiple types of solid tumors.

## Acknowledgements

The authors thank the patients who generously agreed to participate in this study and are grateful for their contribution. We also thank the expertise and assistance of the Dean’s Flow Cytometry and Microscopy Cores at Mount Sinai, Emily Mongeau, Ellen Fraga, Mavis Pancake, and Christopher Bare. This work was funded by NIH R01 CA201189 (N.B.), NIH R01 CA249175 (N.B.), T32 AI007605 (A.M.F.), T32 AI007647 (A.M.F.), and a Translational Team Science Award from the Department of Defense, CA181008 (N.B., J.P.S.).

## Author Contributions

A.M.F. and N.B. conceptualized the study, designed the experiments, and acquired funding . A.M.F. performed experiments, analyzed data and wrote the manuscript. D.Y. designed and performed experiments and analyzed data. M.A.T. designed experiments and analyzed data. S.B., J.H.N., and M.D.G. conceptualized experiments. E.G.G., A.A., A.P. and K.S. performed experiments. J.P.S. provided clinical specimens and acquired funding. F.A, H.A., and J.D. provided clinical specimens. A.H. provided scRNASeq and CyTOF data for HD NK cells. J.T. analyzed CyTOF data. All authors read the final manuscript.

## Declaration of Interests

N.B. is an advisor, consultant, or board member for Apricity, Carisma Therapeutics, CureVac, Genotwin, Novartis, Primevax, Rome Therapeutics, Tempest Therapeutics, and ATP, and has received research support from Dragonfly Therapeutics, Harbour Biomed, and Regeneron. A.H. receives research funding from Astra Zeneca and has served on advisory boards for Purple Biotech and UroGen. M.D.G. receives research funding from Bristol Myers Squibb, Novartis, Dendreon, Astra Zeneca, Merck, and Genentech, and has served as an advisory board member/consultant for Bristol Myers Squibb, Merck, Genentech, AstraZeneca, Pfizer, EMD Serono, SeaGen, Janssen, Numab, Dragonfly, GlaxoSmithKline, Basilea, UroGen, Rappta Therapeutics, Alligator, Silverback, Fujifilm, Curis, Gilead, Bicycle, Asieris, Abbvie, Analog Devices, Veracyte, Daiichi, and Aktis. A.M.F., M.A.T., D.Y., J.H.N., S.B., E.G.G., K.S., A.A. A.P., F.A, H.A., J.T., and J.P.S. declare no competing interests.

## Materials and Methods

### Human specimens

Informed consent was obtained from all patients prior to collection of whole blood, urine, and/or primary tumor tissue under an IRB-approved, genitourinary biorepository protocol at the Tisch Cancer Institute at Mount Sinai. Leukopaks from healthy volunteers were purchased from the New York Blood Center. PBMC were isolated from patient blood via density gradient centrifugation. Briefly, venous blood was collected into EDTA-coated vacutainers (BD Biosciences), diluted 1:2 with DPBS (Gibco), and centrifuged over Ficoll-Paque Plus (GE Healthcare). PBMC were collected and washed with DPBS prior to RBC lysis with ACK (Gibco). Freshly resected tumor tissue was disassociated immediately or placed into tissue storage solution (Miltenyi) at 4° overnight. Tumor tissue was then weighed and dissociated at 37°C with a GentleMACS using program 37C_h_TDK_2 and human tumor dissociation kit reagents (Miltenyi). Cells were passed sequentially through 100µM, 70µM, and 40 µM mesh strainers and washed with DPBS prior to RBC lysis. All NK cell functional assays, scRNASeq, and most immunophenotyping was conducted using fresh cells. Any remaining PBMC or tumor cell suspensions were frozen at -80°C overnight in 90% FBS and 10% DMSO at a rate of -1°C /minute using a Mr. Frostee (Thermo Fisher) prior to storage in vapor phase liquid nitrogen.

### Flow cytometry and antibodies

PBMC and single-cell suspensions derived from primary tumor tissue were stained with a fixable viability dye (Thermo Fisher) on ice for 20’ in DPBS. Cells were washed and F_c_ receptors blocked for 10’ on ice in cold FACS buffer (2% BSA, 1mM EDTA in DPBS). Surface molecules were stained in FACS buffer on ice for 30’ before washing and acquisition. For intracellular staining, cells were additionally fixed, permeabilized, and stained using CytoFix/CytoPerm reagents (BD). Cells were interrogated on LSR Fortessa, LSR Fortessa X-20, or Attune cytometers and FACSDiva v8.0.2 software. Data was analyzed with FlowJo v10.8.2 (BD), and statistics and plots generated using Prism 9.5.0 (GraphPad). Antibodies were purchased from BioLegend, eBiosciences, Miltenyi and BD Biosciences, clones indicated in parantheses: CD45 (HI30), CD3 (SK7), CD14 (63D3), CD19 (HIB19), CD16 (3G8), CD56 (NCAM 16.2), CD4 (RPA-T4), CD8 (RPA-T8), Tim-3 (F38-2E2), TIGIT (MBSA43), PD-1 (EH12.2H7), CD107a (H4A3), Granzyme B (GB11), IFNγ (B27), TNFα (Mab11), NKG2A (REA110), SLAMF6 (NT7), CD4 (RPA-T4), CD8 (RPA-T8), HLA-DR (L243), CD123 (6H6), CD11c (Bu15), Sirpl1 (15-414), and Clec9a (8F9).

### NK cell functional assays

PBMC or single cell suspensions from tumor tissue were obtained as described above. Cells were cultured at 37°C overnight in 96 well, V-bottom plates at a density of .5-1*10^6^ cells/well in complete media (cRPMI) (RPMI containing 10% FBS, 1% penicillin-streptomycin, 1% L-glutamine, and 1% non-essential amino acids) containing 5ng/mL IL-15 (Peprotech) to activate NK cells. The following morning, K562 cells (ATCC) were added to cultures at an effector to target cell ratio of 5:1 in cRPMI containing IL-15 in addition to monensin (2µM) and anti-CD107a (.2µg/mL). After 4h, cells were stained for surface markers, washed, fixed and permeabilized using CytoFix/CytoPerm reagents (BD Biosciences), followed by intracellular staining for IFNγ, TNFα, and Granzyme B (where shown). For *ex vivo* blockade studies, low endotoxin, sodium-azide-free, functional grade anti-Tim-3 (F38-2E2) and/or anti-TIGIT (MBSA34 or 741170) or control IgG1 was added to cultures at 10µg/mL for 1h at 37°C prior to the addition of K562 target cells. For selected experiments, parallel wells without added monensin were used to collect supernatants after NK:K562 co-culture. These supernatants were analyzed for secreted cytokines and cytolytic effector molecules by flow cytometry, using a commercially available bead-based immunoassay (NK/CD8 Legendplex, Biolegend). For Antibody Dependent Cellular Cytotoxicity (ADCC) assays Raji cells (ATCC) were used at a 5:1 E:T ratio in place of K562 target cells, and non-fucosylated anti-CD20 mAb or non-fucosylated control IgG (both InvivoGen) was added at 10ug/mL in the presence of anti-LAMP-1 for 4 hours, prior to staining for flow.

### Tumor rest assays

Single cell suspensions from tumor tissue were isolated as described above and plated in cRPMI containing either 50U/mL IL-2 or 1ng/mL IL-15 (Peprotech) in 12-well culture plates. On day 2, cells were centrifuged, and the media replaced with fresh cRPMI and IL-2 or IL-15. On day 4, cells were centrifuged, and media replaced with fresh cRPMI containing 5ng/mL rhIL-15 to activate NK cells. The following morning K562 cells were added to cultures to assess NK function as described above.

### Glucose uptake assays

To measure the frequency of cells able to uptake glucose, PBMC or cells from dissociated tumor were incubated in Krebs-Ringers-HEPES (KRH) for 15 min at 37°C, washed with KRH, then resuspended in KRH containing 100uM of 2-NBDG, a fluorescent glucose analog (ThermoFisher). For 10 min at 37°C. Cells were subsequently washed and stained with lineage markers to identify NK and T cells, and the frequency of 2-NBDG^+^ cells determined by flow cytometry.

### Hypoxia assays

Hypoxic cell culture (4% O_2_) was performed by enclosing the cell culture plate in a plastic pouch with an O_2_ scavenger pack and an O_2_ meter (nBionix hypoxic cell culture kit, Bulldog Bio). After the concentration inside the pouch fell to 4% O_2,_ the cell culture plate was clamped off from the O_2_ scavenger pack and was placed inside a standard incubator for the duration of the experiment (3-4d).

### Phospho-protein arrays

The proteome profiler human phosphoimunoreceptor array kit (R&D Systems) was used to determine the phosphorylation status of the ITSM of SLAMF6, according to the manufacturer’s protocol. Briefly, cells were cultured under the conditions described, then harvested to make protein lysates. Total protein concentration was measured using a standard Bradford assay (Pierce Detergent Compatible Bradford Assay Kit, ThermoFisher), and 150ugs of total protein was incubated on membranes pre-spotted with capture antibodies for 59 immune-related receptors. Lysates were incubated with membranes overnight at 4°, washed, and probed with an anti-phosphotyrosine-HRP detection antibody before standard chemiluminescent detection on film. Films were semi-quantified with QuickSpots software (Ideal Eyes https://idealeyes.com/products/QuickSpots.php).

### O-link

The Olink Immuno-Oncology panel was used to measure the amounts of 92 soluble proteins present in healthy or BCa patient plasma and BCa patient urine. Plasma was collected by centrifuging whole blood at 1,000g for 10’ at 4°C and urine was centrifuged at 500g for 5’ at 4°C to remove any cells present. Each analyte was detected using a pair of antibodies, each containing complimentary oligonucleotides, with quantification by quantitative reverse transcriptase quantitative polymerase chain reaction (RT-qPCR). 1 μL of plasma or cell-free urine was added to 3 μL of an O-Link binding buffer and incubated overnight at 4°C. The following morning an O-link PCR master mix was added to each well and pre-amplified. For detection, 2.8 μL of pre-amplified product from each well was mixed with 7.2 μL Olink detection mix, transferred to an Olink fluidic circuit, and run on a BioMark qPCR instrument (Standard BioTools). Samples were run in singlet with recommended technical controls. Detailed protocols are available on the Olink website (https://www.olink.com). QC and normalization were done by Olink guidelines. Data was analyzed using the ΔΔCt method and an Olink normalized protein expression manager, and read out on a log2 scale as NPX (normalized protein expression),

### CyTOF on HD PBMC

Frozen HD PBMC (n=20) were obtained from an IRB approved protocol at the Karolinska Institute. PBMC were thawed and recovered in complete RPMI overnight without exogenous cytokines before being stimulated with either 1 ng/mL of IL-15 (Peprotech) or 1 ng/mL of IL-15+ 5 ng/mL TGF-β (Peprotech) for 6 days, with media being replaced with fresh media containing the appropriate cytokines every 2 days. After the 6-day culture PBMC were transferred to V-bottom plates and co-cultured with K562 cells (ATCC) (E/T ratio, 10:1) for 4 hours at 37°C in the presence of 10ug/mL brefeldin A and .5ug/mL of anti-LAMP-1 mAb. 15 minutes prior to the end of K652 co-culture the media was replaced with media containing a 1:500 dilution of rhodamine viability dye (Standard BioTools) for subsequent analysis by CyTOF. For ADCC assays PBMC were cultured as above, but Raji cells with bound anti-human CD20-IgG1 mAb (BioLegend, clone QA20A02) were used as targets. After K562 or Raji co-culture HD PBMC were washed twice with CyFACS buffer (DPBS containing 2mM EDTA and 0.1% BSA), and cells from each donor were barcoded by staining with anti-human CD45 mAb conjugated to 195Pt, 196Pt, 198Pt (clone HI30, Biolegend) or anti-human B2m mAb conjugated to 195Pt, 196Pt, 198Pt (clone 2M2, Biolegend) antibodies. Samples were then washed with CyFACS buffer, fixed with 1% PFA, resuspended in BD FACS Lyse solution, and stored at -80°C until staining for intracellular proteins. The cells were thawed, washed in CyFACS buffer, and centrifuged at 300g for 15 minutes at 4°C. Samples were then permeabilized with Maxpar Barcode permeabilization buffer (Standard BioTools) and intracellular barcoding performed by staining with anti-human CD45 mAb-conjugated to Pd104, Pd105, Pd106, Pd108 and Pd110 (Standard BioTools) for 30min at R.T. Samples were then pooled, resuspended in permeabilization buffer (eBioscience) and stained for intracellular targets using the panel described in **Table S2** for 30min at 4°C. Finally, samples were washed, then incubated with intercalators conjugated to Ir191/Ir193 (Standard BioTools) diluted 1:4,000 in PBS containing 1% formaldehyde for 30 minutes at R.T. Samples were washed a final time then resuspended in CSM-Ir and kept at 4°C until acquisition on a Helios mass cytometer (Standard BioTools).

### Analysis of CyTOF NK clusters with Depeche

For the clustering analysis, one donor with fewer than 100 events in both TGF-stimulated and non-stimulated samples was excluded. For the remaining samples, either all events, (for samples with <500 total events) or 500 random events (for samples >500 total events) were used for clustering, so as to minimize the influence of samples containing high cell numbers. For clustering, Determination of Essential Phenotypic Elements of Cells in High-dimensional Entities” (DEPECHE) was used^85^. This is a penalized, k-means-based method that automatically identifies the optimal cluster resolution by iterative clustering with different penalties and Adjusted Rand Index (ARI) comparisons between the results. It also penalizes non-informative markers on a per-cluster basis. With this method, 4 clusters were identified as giving rise to an optimally reproducible and statistically interpretable model. After this clustering step, all previously excluded cells were assigned to the 4 identified clusters so as to enable the highest possible statistical power in subsequent analyses.

### Immunohistochemistry

#### Tim-3, PD-1, and CD3

Serial 5µM FFPE sections were stained for CD3 (Clone D7A6E, Cell Signaling Technology, 1ug/mL), Tim-3 (Clone D5D5RTM, Cell Signaling Technology, 2ug/mL) and PD-1 (Clone NAT-105, Sigma 2ug/mL), with the first serial section reserved for H&E staining. Slides were deparaffinized and rehydrated, followed by staining of CD3 and PD-1 w with a BenchMark Ultra autostainer (Roche). Tim-3 was stained manually after heat-induced epitope retrieval using the “high pressure” program on an electric pressure cooker (Cuisinart CPC-600) for 7 minutes in Tris-EDTA buffer (pH 9.0). The primary Ab was incubated O/N at 4°. All targets were detected with HRP-conjugated 2°Abs directed against the host species of each 1° Ab, followed by chromogenic revelation with 3, 3-diaminobenzadine. Images were acquired under brightfield illumination on a Nikon Eclipse Ti-s inverted microscope or a Pannoramic Flash 250 digital slide scanner (3D Histech).

#### HIF-1l1

A commercially available human multiple tumor tissue microarray comprised of 6µM FFPE sections of kidney, ureter, bladder, and stomach tumors, as well as unmatched normal tissue controls was used for HIF-1l1 IHC (Novus Biologicals). A rabbit anti-human HIF-1l1 monoclonal Ab was used (Clone 2443C, Novus, 1µg/mL). Slides were deparaffinized, rehydrated, and subjected to heat-induced epitope retrieval using the “simmer” program on an electric pressure cooker (Cuisinart CPC-600) for 20 minutes in High pH antigen retrieval solution (eBioscience). Primary Abs were incubated O/N at 4°and detected with an HRP-conjugated 2°Ab (Vector Laboratories). Chromogenic revelation was performed with 3, 3-diaminobenzadine (HIF-1l1) and Vector Red (Nkp46). Images were acquired under brightfield illumination on a motorized Zeiss Axioimager to acquire tiled images. The multiple tumor tissue array was obtained from Novus.

### Sample preparation for scRNASeq

BCa patients, live, CD45^+^ singlets were FACS-sorted (BD FACSAria II, 100µM nozzle, 4° C) from either Ficoll-isolated PBMC or freshly dissociated tumor tissue. Cells were sorted into DPBS + .04% BSA, resuspended at 1000 cells/µL, and loaded onto a Chromium controller (10x Genomics) for encapsulation into droplets, lysis, barcoding, reverse transcription, and library preparation using Single Cell 3’ v2 reagents. For HD NK cells, PBMC were isolated as above, followed by depletion of non-NK lineages using immuno-magnetic beads (EasySep human NK cell enrichment kit, Stemcell Technologies). Samples were sequenced in dedicated flow cells on a HiSeq 2500 (Illumina) to an average depth of 91,787, 76,918, and 71,692 reads/cell for HD NK cells, BCa patient PBMC, and TIL respectively.

### Pre-processing and analysis of scRNASeq

Raw Illumina sequencing data was converted to FASTQ, demultiplexed and aligned to the GRCh38 reference genome using CellRanger v3.0.0 (10x Genomics). Subsequent analysis was performed using Seurat v5^54^ (https://satijalab.org/seurat) in RStudio v2021.09.0. For QC, cells expressing more than 2,500, and fewer than 300 unique genes, as well as cells containing more than 10% mitochondrial transcripts, were discarded. Genes expressed in fewer than 5 cells were also discarded. Expression data was then normalized by a factor of 10,000 and log-transformed. Next, highly variable genes in each sample were identified by binning the average expression and dispersion of each gene using Seurat’s FindVariableFeatures function. To enable direct comparisons between NK cells from different tissues, disease stages, and donors, the resulting matrices from each donor of HD PBMC NK cells (n=5), BCa PBMC (n=7), and TIL (n=6) were aligned and merged using the FindIntegrationAnchors/IntegrateData approach in Seurat. Principal component analysis (PCA) was run on the scaled data, and dimensionality reduction using UMAP was performed on the first 20 principal components. Next, graph-based clustering was performed, with iterative adjustment of the resolution parameter in the FindClusters command to maximize cluster membership amongst donors. Clusters were visualized using UMAP. Differentially expressed genes (DEG) were identified using a non-parametric Wilcoxon rank sum test (Seurat FindMarkers and FindAllMarkers) on genes expressed by ≥ 5% of cells with a ≥ 5% expression difference across the clusters being tested.

10,345 cells from TIL that were annotated as tumor, epithelial, stromal, or endothelial cells were removed, resulting in a matrix comprised of 68,192 cells (7,812 HD NK cells, 39,963 PBMC, and 20,241 TIL) across 19 clusters at resolution .8. Of these, 18,762 cells in clusters 3, 4, 10, and 13 were annotated as NK cells by comparing the module scores of published NK cell gene signatures ^26,55,57,125^, as well as by expression of additional, canonical NK cell transcripts. These NK cell clusters were then sub-clustered at resolution 0.2 to identify DEG across tissue and disease. Pathway enrichment analysis was performed taking as input the top 100 significant DEG (by adjusted p-value) for NK cells from each tissue as input. Pathways were queried using Enrichr ^126^ (http://amp.pharm.mssm.edu/Enrichr) and visualized with GOPlot ^127^ and Prism v8. Gene signature scores were calculated by subtracting the expression values of a randomly selected set of control genes from that of the genes comprising each signature using the AddModuleScore function in Seurat ^128^. RNA pseudotime analysis was performed using Monocle v2.12 ^77^. Briefly, NK cell genes that were significantly differentially expressed across tissue and cluster of origin were used to order cells in Monocle. Dimensionality reduction was performed using a max.components parameter of 4.

### SLAMF6 CRISPR in NK-92 cells

NK-92 cells were cultured in Myelocult H5100 (StemCell Technologies) supplemented with 100U/mL IL-2 (Peprotech). Low passage cells not exceeding 3*10^5^/mL were used for deletion of SLAMF6. Briefly, SLAMF6 tracrRNA and crRNA (Alt-R CRISPR-Cas9 crRNA: 5’-/AI+R1/rGrGrU rUrUrC rArUrG rGrGrG rUrArC rUrArU rGrArG rUrUrU rUrArG rArGrC rUrArUrGrCrU /AltR2/ -3’, IDT) were annealed to generate gRNA targeting SLAMF6. gRNA was incubated with Cas9 nuclease v3 (IDT) and electroporation enhancer (IDT) to generate a Ribonucleoprotein (RNP). 5*10^5^ NK-92 cells were resuspended in Solution 2 + Mannitol (5 mM KCl, 15 mM MgCl_2_, 15 mM HEPES, 150 mM Na_2_HPO_4_ /NaH_2_ containing 50 mM mannitol, pH 7.2)^129^ and the RNP was added. Nucleofections were performed in a volume of 20uL in cuvettes (Lonza) with Pulse Code CM-137 on an Amaxa 4D-Nucloefector X (Lonza). Cells were recovered and expanded in NK MACS media (Miltenyi) supplemented with 500U/mL IL-2. On d5-d7 SLAMF6^-^ NK-92 were sorted on a FACSAria (BD Biosciences) and allowed to expand again for 7-14 days prior to co-culture with K562 to assess function. Typical d5-d7 deletion efficiency was ∼70% and typical sort purity was >90%.

### Bulk RNAseq of SLAM6-KO and Hypoxic NK-92 cells

Total RNA was extracted from 1*10^6^ WT or sorted SLAMF6-KO NK-92 cells using an RNA Micro Kit (Zymo Research). After PolyA selection libraries were prepared and lllumina sequencing of 20-30 million paired-end reads performed. DESeq2 was used to identify DEG between WT and SLAMF6-KO cells. 2,024 significantly DEG were identified with significance defined as a gene having an adjusted p-value of < 0.05 and an absolute log2 fold change > 1 as determined by the Wald test. Samples were sequenced and DEG determined by Genewiz, and volcano plots were generated using the R package EnhancedVolcano (https://github.com/kevinblighe/EnhancedVolcano).

### SLAMF6 and clinical outcomes

Kmplot.com was used to query the association between SLAMF6 expression and BCa patient PFS, as well as ICB response across 13 cancers <u>(www.Kmplot.com).</u> Kmplot uses a manually-curated dataset with expression and survival data being obtained from TCGA, GEO, and EGA. The queried clinical cohort is divided into two groups based on expression of the queried gene. A Kaplan-Meier plot is generated and the hazard ratio, 95% confidence interval, and logrank p-values are computed^130^.

## Notes

### Summary of Updates

Added a co-author and therefore updated the author list.

