## Supplementary material for "Natural Killer Cell Dysfunction In Human Bladder Cancer Is Caused By Tissue-Specific Suppression of SLAMF6 Signaling": Manuscript and Figures (from another version of this preprint: version 1, 2024-05-03)

**Table S1: Clinical characteristics of patient cohort**

| Clinical State at Collection | Total (n=68) |
|---|---|
| NMIBC | 39 (57.4%) |
| MIBC | 25 (36.8%) |
| Metastatic | 4 (5.9%) |
| Tumor Stage | |
| pTis | 7 (10.3%) |
| pTa | 20 (29.4%) |
| pT1 | 17 (25%) |
| pT2 | 10 (6.8%) |
| pT3 | 5 (7.3%) |
| pT4 | 4 (5.9%) |
| No Stage | 5 (7.3%) |
| pT0 | 2 (2.9%) |
| Grade | |
| Low | 10 (6.8%) |
| High | 29 (42.6%) |
| Previous Treatment | |
| None | 46 (67.6%) |
| Chemotherapy | 13 (19.1%) |
| Radiation | 0 (0%) |
| Immune Checkpoint Blockade | 2 (2.9%) |
| BCG | 9 (13.2%) |
| Surgical Procedure | |
| TURBT | 52 (76.5%) |
| Cystectomy | 11 (16.2%) |

#### Table S2: CyTOF Antibody Panel

| Metal | Metal Source | Target | Antibody Clone | Antibody Source |
|---|---|---|---|---|
| 89 Y | Standard BioTools | CD57 | HCD57 | BioLegend |
| 113 In | Standard BioTools | CD45 | H130 | BioLegend |
| 115 In | Standard BioTools | HLA-DR | L243 | BioLegend |
| 141 Pr | Standard BioTools | KIR3DL1 | DX9 | R&D Systems |
| 142 Nd | Standard BioTools | KIR2DL3 | REA147 | Miltenyi |
| 143 Nd | Standard BioTools | CD94 | HP-3D9 | BD |
| 144 Nd | Standard BioTools | IFNγ | B27 | BioLegend |
| 145 Nd | Standard BioTools | CD4 | RPA-T4 | BioLegend |
| 146 Nd | Standard BioTools | CD8α | RPA-T8 | BioLegend |
| 147 Sm | Standard BioTools | Ki-67 | B56 | BD |
| 148 Nd | Standard BioTools | CD16 | 3G8 | BioLegend |
| 149 Sm | Standard BioTools | CEACAM-1 | 5F4 | Dr. R. Blumberg |
| 150 Nd | Standard BioTools | PLZF | 6318100 | R&D Systems |
| 151 Eu | Standard BioTools | KIR2DL1 | REA284 | Miltenyi |
| 152 Sm | Standard BioTools | KIR2DL2/L3/S2 | REA1007 | Miltenyi |
| 153 Eu | Standard BioTools | PD-1 | EH12.2H7 | BioLegend |
| 154 Sm | Standard BioTools | FcERIγ | Polyclonal | Millipore |
| 155 Gd | Standard BioTools | NKG2D | 1D11 | BioLegend |
| 156 Gd | Standard BioTools | CD14 | HCD14 | DVS Science |
| 158 Gd | Standard BioTools | Granzyme B | REA226 | Miltenyi |
| 159 Tb | Standard BioTools | B7-H3 | 185504 | R&D Systems |
| 160 Gd | Standard BioTools | Siglec-7 | 194211 | R&D Systems |
| 161 Dy | Standard BioTools | TIM-3 | 344823 | R&D Systems |
| 162 Dy | Standard BioTools | NKp46 | BAB281 | DVS Science |
| 163 Dy | Standard BioTools | LILRB1 | HP-F1 | eBioscience |
| 164 Dy | Standard BioTools | TIGIT | MBSA43 | eBioscience |
| 165 Ho | Standard BioTools | VISTA | 730804 | R&D Systems |
| 166 Er | Standard BioTools | NKG2C | 134591 | R&D Systems |
| 167 Er | Standard BioTools | DNAM-1 | 11A8 | BioLegend |
| 168 Er | Standard BioTools | CD3 | UCHT1 | BioLegend |
| 169 Tm | Standard BioTools | NKG2A | 3169013B | DVS Science |
| 170 Er | Standard BioTools | X-Biotin | 1D4-C5 | BioLegend |
| 171 Yb | Standard BioTools | KIR2DL1/S1 | REA1010 | Miltenyi |
| 172 Yb | Standard BioTools | CD132 | TUGH4 | BioLegend |
| 173 Yb | Standard BioTools | CD25 | BC96 | BioLegend |
| 174 Yb | Standard BioTools | CD122 | TU27 | BioLegend |
| 175 Lu | Standard BioTools | LAG-3 | 874501 | R&D Systems |
| 176 Yb | Standard BioTools | CD56 | HCD56 | BioLegend |
| | | Biotinylated-LAMP-1 | H4A3H4A3 | BioLegend |

**Table S3: Clinical characteristics of single-cell RNAseq cohort**

| Patient ID | Disease Stage | Pathology | Surgical Procedure | Tx History | Age | Sex | Tissues Analyzed | Isolation Method |
|---|---|---|---|---|---|---|---|---|
| BC 155 | MI | pT2 | TURBT | Unknown | 59 | M | PBMC+TIL | Sorted $CD45^{+}$ |
| BC 158 | MI | pT3 | TURBT | None | 66 | M | PBMC+TIL | Sorted $CD45^{+}$ |
| BC 166 | NMI | pTa | TURBT | None | 83 | F | PBMC+TIL | Sorted $CD45^{+}$ |
| BC 167 | MI | pT4b | TURBT | Ipimilumab+Nivolumab | 67 | M | PBMC+TIL | Sorted $CD45^{+}$ |
| BC 168 | MI | pT3b | Cystectomy | None | 82 | M | PBMC+TIL | Sorted $CD45^{+}$ |
| BC 170 | NMI | pT1 | TURBT | BCG | 86 | M | PBMC+TIL | Sorted $CD45^{+}$ |
| BC 183 | MI | pT2a | Cystectomy | BCG | 72 | M | PBMC | Sorted $CD45^{+}$ |
| HD 205 | HD | N/A | N/A | N/A | Unknown | Unknown | PBMC | CD56 negative selection |
| HD 206 | HD | N/A | N/A | N/A | Unknown | Unknown | PBMC | CD56 negative selection |
| HD 207 | HD | N/A | N/A | N/A | Unknown | Unknown | PBMC | CD56 negative selection |
| HD 208 | HD | N/A | N/A | N/A | Unknown | Unknown | PBMC | CD56 negative selection |
| HD 210 | HD | N/A | N/A | N/A | Unknown | Unknown | PBMC | CD56 negative selection |

#### Table S4: Gene signatures used for scRNAseq

| Gene Signature | Genes in Signature | # Genes in Signature | Source | PMID |
|---|---|---|---|---|
| **Human NK Cells** | CD160, CD244, CHST12, CST7, GNLY, IL18RAP, IL2RB, KLRC1, KLRC3, KLRD1, KLRF1, PRF1, XCL2 | 13 | Crinier et al, Immunity 2018 | 30413361 |
| **Human NK Cells** | GNLY, KLRC3, KLRD1, KLRF1, NCR1 | 5 | Barry et al, Nature Medicine 2018 | 29942093 |
| **Human NK Cells_Resting** | PRF1, GNLY, IL2RB, CCL5, CD247, GZMB, NKG7, GZMA, KLRB1, GZMH, CST7, TRDC, KLRF1, KLRD1, CTSW, APOBEC3G, CD2, IL18RAP, SELL, CD300A, ZAP70, PVRIG, TRBC1, TBX21, GZMM, LCK, PTPRCAP, DEAF4, ITK, PTGER2, TTC38, FAIM3, CD160, GPR65, CD37, SKAP1, GZMK, ADRB2, CD244, CD69, LAT, KLRG1, CD7, CAMP, CD96, PTGDR | 48 | Newman et al, Nature Methods 2015 | 25822800 |
| **Human NK Cells_Activated** | PRF1, GNLY, IL2RB, CCL5, CD247, GZMB, NKG7, GZMA, KLRB1, GZMH, CST7, TRDC, KLRF1, KLRD1, CTSW, APOBEC3G, CD2, IL18RAP, LTB, DEAF4, CD300A, ZAP70, PVRIG, TRBC1, TBX21, GZMM, LCK, PTPRCAP, CCND2, ITK, PTGER2, NCR3, KIR2DS4, LTA, SAMSN1, IRF8, FASLG, TNFSF14, GPR18, TXK GPR65, ADRB2, CD244, CD69, CD7, CD96, PTGDR | 48 | Newman et al, Nature Methods 2015 | 25822800 |
| **Human NK Cells** | Please see PMID 30110645 | 500 | Balan et al, Cell Reports 2018 | 30110645 |
| **NK Tissue Residency** | CD69, CXCR6, ITGA1, ITGAE, RGS1 | 5 | Dogra et al, Cell 2020<br>Tang et al, Cell 2023 | 32059780<br>37607536 |
| **Human CD56$^{bright}$ NK Cells** | Please see PMID 32019778 | 120 | Ercolano et al, Cancer Immunology Research 2020 | 32019778 |
| **TGF-β Signaling** | ACTA2, ACTG2, ADAM12, ADAM19, CNN1, COL4A1, CTGF, CTPS1, FAM101B, FSTL3, HSPB1, IGFBP3, PXDC1, SEMA7A, SH3PXD2A, TAGLN, TGFB1, TNS1, TPM1 | 19 | Mariathasan et al, Nature 2018 | 29443960 |
| **Hypoxia** | AL2, ALDOA, AM, BCL2, BCL2L1, FOS, JUN, SRC, CBP, CCR6, CEBPB, ENOLAS, FAS, FASLG, FKBP52, GAPDH, GLUT1, GLUT3, CSF2, IFNG, IFNB1, IL13, IL1R1, IL2RA, IL4, IL5, IL6R, IL8, IL10R, IL15R, ITGA6, ITK, JUND, CLB54, MAPP3K5, MAP2K1, MAP2K2, MIF, NFATC1, NFKB2, P53, TGM6, TNF, TNFRSF1B, TRADD, TRAIL, TRAP1, VEGF | 48 | Azizi et al, Cell 2018<br>Makino et al, JI 2003<br>Benita et al, NAR 2009 | 29961579<br>14662854<br>19491311 |
| **Glucose Deprivation** | NRN1, FAM129A, 1L23A, SPRY1, GADD45B, HSPA1A/B, GPNMB, HAVCR2, SOAT2, ADAMTS6 | 10 | Azizi et al, Cell 2018<br>Ho et al, Cell 2015 | 29961579<br>26321681 |
| **Cytoxicity** | EOMES, TBX21, GZMB, PRF1, FASL, GZMH, GZMA, GNLY, GZMK, GZMM, FCGR3A | 11 | Azizi et al, Cell 2018<br>Glimcher et al, NRI 2004<br>This paper | 29961579<br>15516969 |
| **Cytokines** | IFNG, TNF, CCL5, XCL1/2, FLT3LG | 6 | This paper | |
| **Glycolysis** | ALDOA, ALDOB, ALDOC, BPGM, ENO1, ENO2, GALM, GCK, GPI, HK2, HK3, PFKL, PGAM2, PGK1, PGK2, PGM1, PGM2, PGM3, PKLR, TPI1 | 20 | Azizi et al, Cell 2018<br>Moreno-Sanchez et al, Biofactors 2019 | 29961579<br>19449450 |
| **Combined Inhibitory SLAMF6 Signaling, Hypoxia, and TGF-β Signature** | HIF1A, ZNF683, INPP5D, SLAMF6, PTPN6, ACTA2, ACTG2, ADAM12, ADAM19, CNN1, COL4A1, CTGF, CTPS1, FAM101B, FSTL3, HSPB1, IGFBP3, PXDC1, SEMA7A, SH3PXD2A, TAGLN, TGFB1, TNS1, TPM1 | 24 | This paper<br>Mariathasan et al, Nature 2018<br>Li et al, Clin Transl Med 2023 | 29443960<br>36245253 |

**Figure S1**

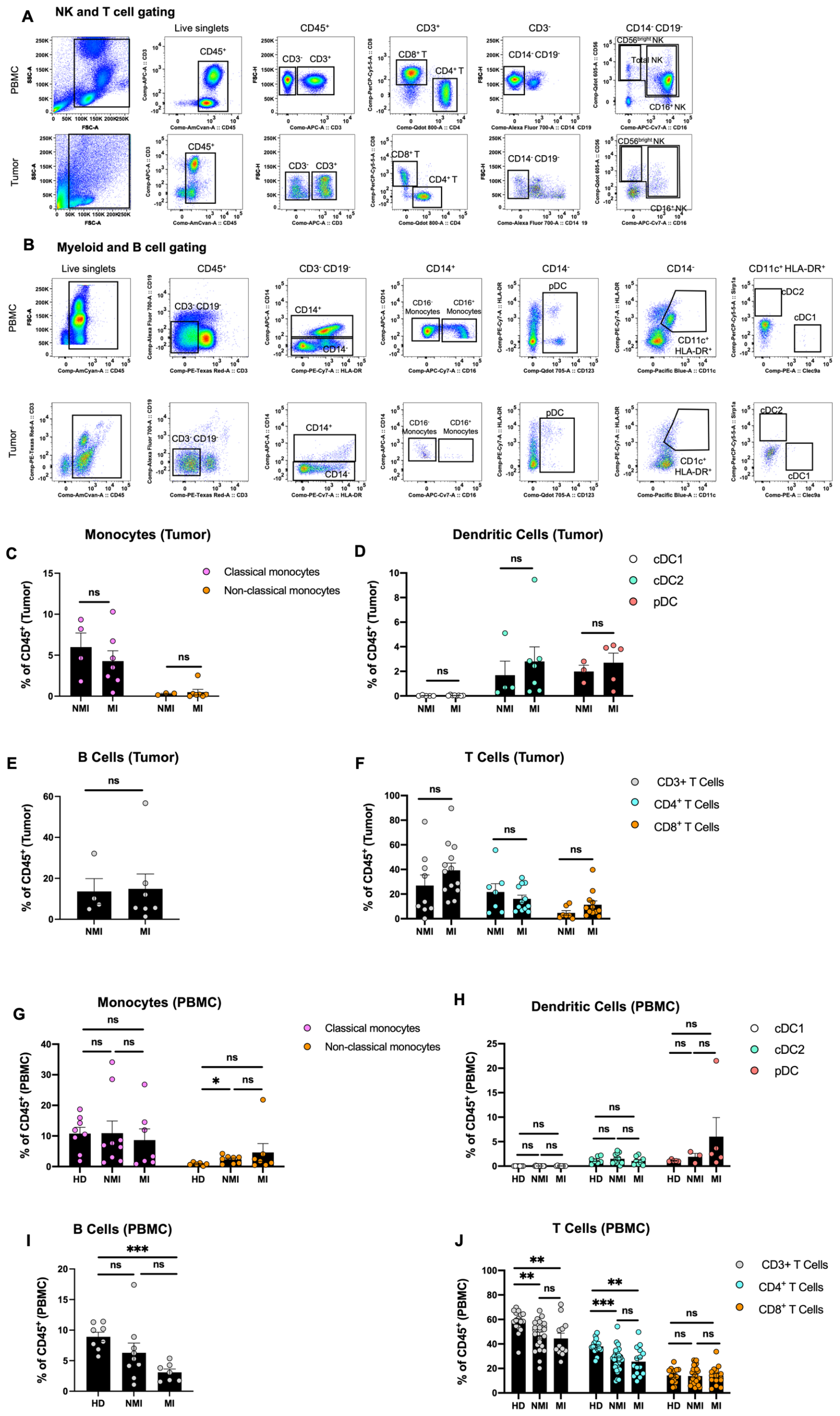

A
NK and T cell gating
Live singlets
CD45+
CD3+
CD3-
CD14- CD19-
PBMC
Tumor
CD3- CD3+
CD8+ T
CD4+ T
CD56bright NK
Total NK
CD16+ NK
B
Myeloid and B cell gating
CD3- CD19-
CD14+
CD14-
CD11c+ HLA-DR+
CD16- Monocytes
CD16+ Monocytes
pDC
cDC2
cDC1
C
Monocytes (Tumor)
Classical monocytes
Non-classical monocytes
% of CD45+ (Tumor)
NMI
MI
ns
D
Dendritic Cells (Tumor)
cDC1
cDC2
pDC
E
B Cells (Tumor)
F
T Cells (Tumor)
CD3+ T Cells
CD4+ T Cells
CD8+ T Cells
G
Monocytes (PBMC)
% of CD45+ (PBMC)
HD
H
Dendritic Cells (PBMC)
I
B Cells (PBMC)
J
T Cells (PBMC)

**Figure S1: Immune composition of bladder tumors and BCa PBMC (A)** Gating strategy to identify NK cells ($CD45^+CD3^-CD14^-CD19^-CD56^+CD16^{+/-}$) and T cells ($CD45^+CD3^+CD4^+CD8^-$ and $CD45^+CD3^+CD4^-CD8^+$) in PBMC and tumor tissue. **(B)** A separate FACS panel was used in a subset of the cohort to identify myeloid lineages and B cells as follows: cDC1 ($CD45^+CD3^-CD14^-CD19^-$HLA-DR+CD11c+Sirp1$\alpha^-$Clecl9a$^+$), cDC2 ($CD45^+CD3^-CD14^-CD19^-$HLA-DR+CD11c+Sirp1$\alpha^+$Clecl9a$^-$), pDC ($CD45^+CD3^-CD14^-CD19^-$CD123+HLA-DR$^{+/-}$), classical and non-classical monocytes ($CD45^+CD3^-CD19^-CD14^+CD16^{+/-}$), and B cells ($CD45^+CD3^-CD19^+$) in PBMC**)** and tumor tissue. Parental gates are indicated above plots. In tumor, isotype controls for CD45 and CD3 were used to determine appropriate placement of gates. **(C)** Frequency of classical and non-classical monocytes **(D),** cDC1, cDC2 ,and pDC, **(E)** B cells, and **(F)** T cells in the TIL of BlCa patients (NMI: n=5, MI: n=8). **(G-I)** Frequency of the same populations in the PBMC of BlCa patients and HD (HD: n=8, NMI: n=9, MI: n=7). HD = Healthy Donor, NMI =Non-Muscle-Invasive, MI=Muscle-Invasive. ns, $p \geq .05$, $*p < .05$, $**p < .01$, $***p < .001$ as determined by unpaired t-test. Error bars show SEM.

### Figure S2

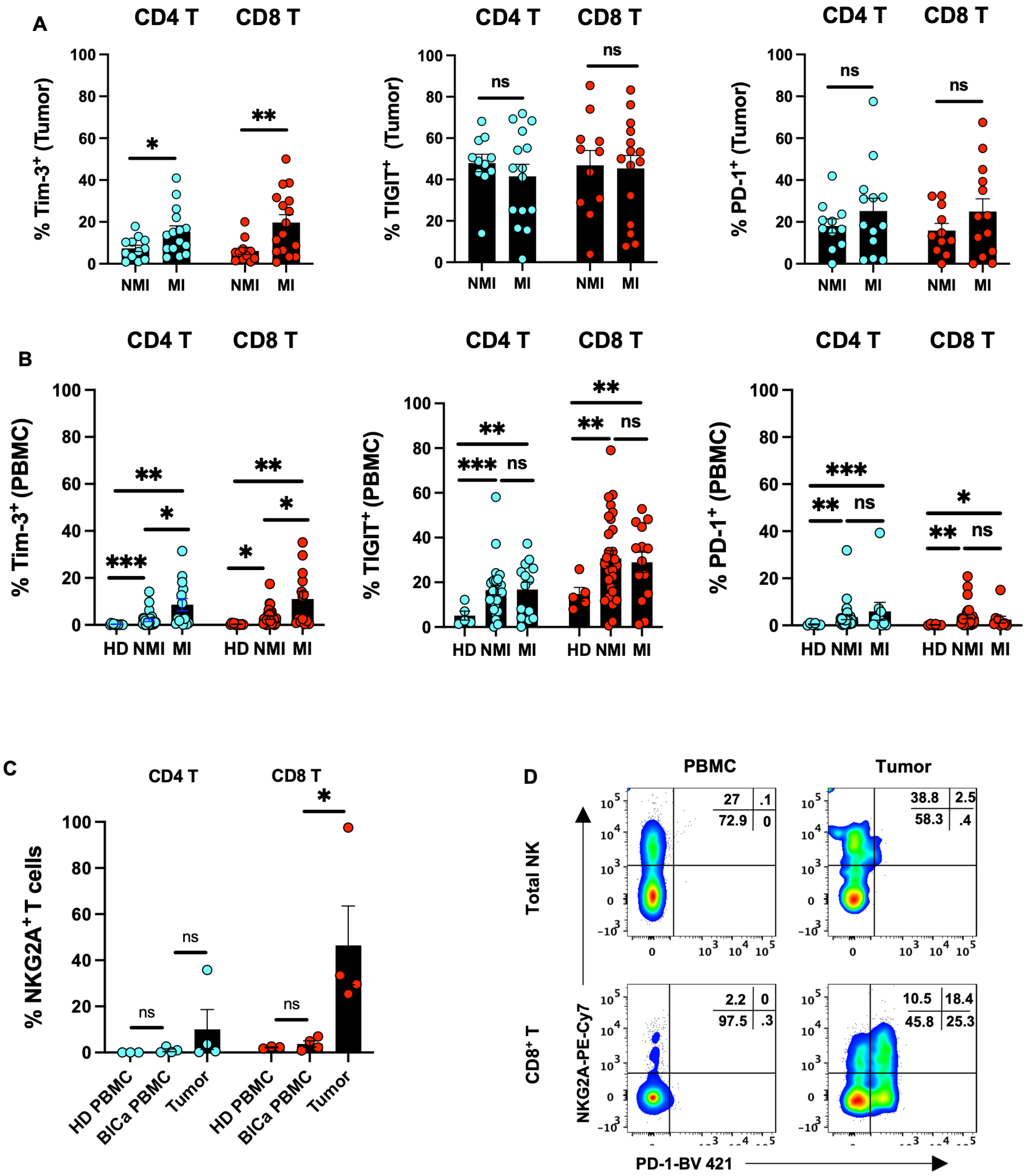

A
CD4 T
CD8 T
% Tim-3+ (Tumor)
% TIGIT+ (Tumor)
% PD-1+ (Tumor)
NMI
MI
ns
B
% Tim-3+ (PBMC)
% TIGIT+ (PBMC)
% PD-1+ (PBMC)
HD
C
% NKG2A+ T cells
HD PBMC
BlCa PBMC
Tumor
D
PBMC
Tumor
Total NK
CD8+ T
NKG2A-PE-Cy7
PD-1-BV 421
27
.1
72.9
0
38.8
2.5
58.3
.4
2.2
0
97.5
.3
10.5
18.4
45.8
25.3

**Figure S2: Bladder cancer imprints conserved disease, stage, and tissue-specific phenotypes on T cells.** PBMC and cell suspensions obtained from patient-matched tumor tissue were analyzed by flow cytometry to determine expression of inhibitory receptors on T cells. **(A)** Frequency of Tim-3$^+$, TIGIT$^+$, or PD-1$^+$ CD4$^+$ and CD8$^+$ T cells in bladder tumors (NMI: n=12, MI: n=14). **(B)** Frequency of Tim-3$^+$, TIGIT$^+$, or PD-1$^+$ CD4$^+$ and CD8$^+$ T cells in BCa patient PBMC (HD: n= 12, NMI: n=29, MI: n=14). **(C)** Frequency of NKG2A$^+$ CD4$^+$ and CD8$^+$ T cells in PBMC (HD: n=3, BCa: n=4), and tumor (n=4). **(D)** Representative FACS plots of NKG2A and PD-1 staining on NK and CD8$^+$ T cells from in the PBMC or tumor of a BCa patient. HD = Healthy Donor, NMI =Non-Muscle-Invasive, MI=Muscle-Invasive. ns, p≥.05, *p<.05, **p <.01, ***p <.001 as determined by unpaired t-test. Error bars show SEM.

### Figure S3

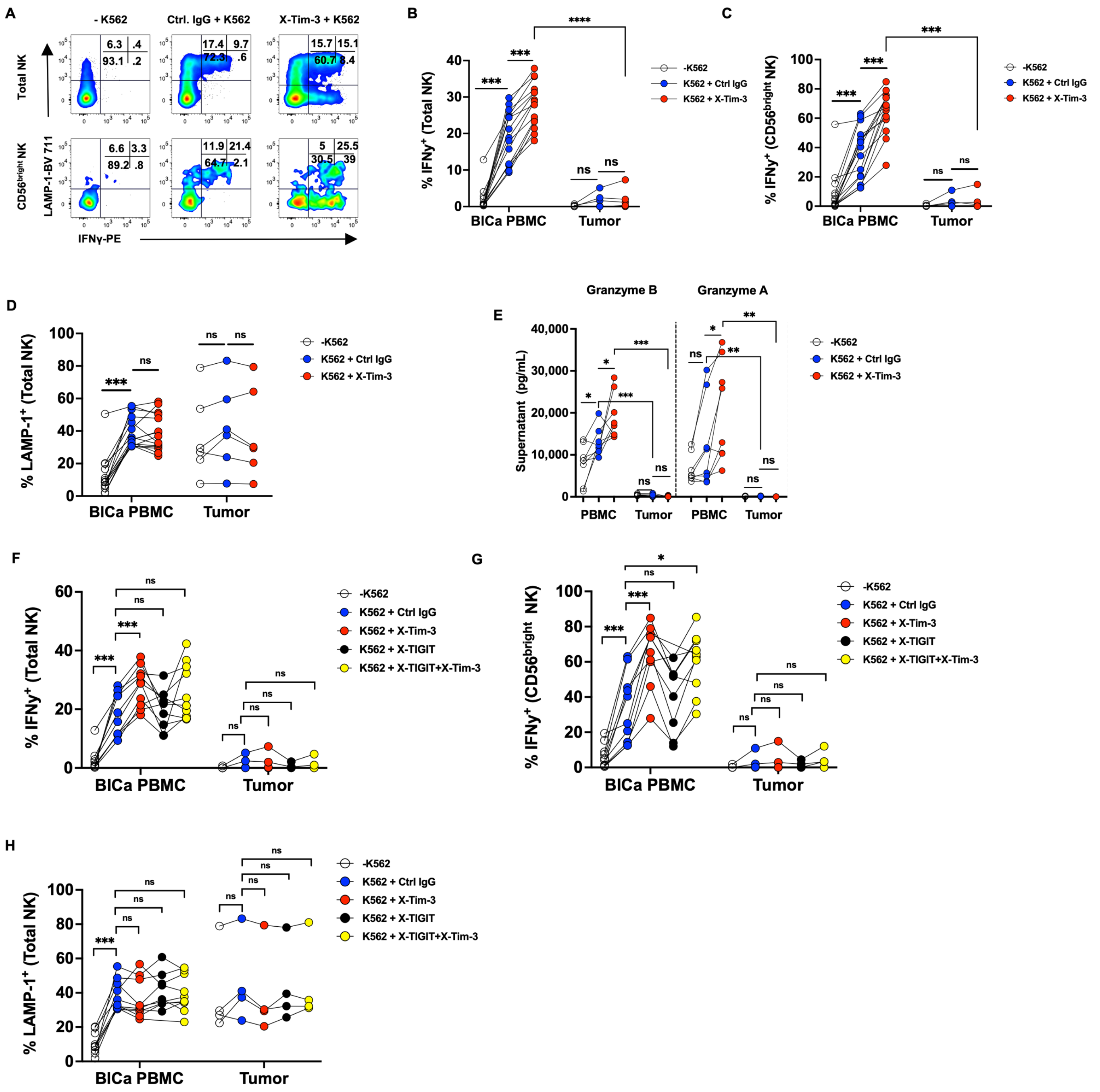

A
B
C
D
E
F
G
H
- K562
Ctrl. IgG + K562
X-Tim-3 + K562
Total NK
CD56bright NK
LAMP-1-BV 711
IFNγ-PE
% IFNγ+ (Total NK)
% IFNγ+ (CD56bright NK)
% LAMP-1+ (Total NK)
BlCa PBMC
Tumor
-K562
K562 + Ctrl IgG
K562 + X-Tim-3
K562 + X-TIGIT
K562 + X-TIGIT+X-Tim-3
Granzyme B
Granzyme A
Supernatant (pg/mL)
PBMC

**Figure S3: *Ex vivo* blockade of Tim-3 enhances the effector function of BCa pbNK cells, while TIGIT blockade does not.** PBMC or cell suspensions from dissociated tumor tissue were activated overnight with 5ng/mL of IL-15. Blocking Abs against Tim-3 (F38-2E2) or ctrl. IgG were added at 10ug/mL for 1h prior to the addition of K562 target cells to elicit IFNγ and degranulation. **(A)** Representative FACS plots for IFNγ and LAMP-1 gated on total NK cells (top) and $CD56^{bright}$ NK cells (bottom) from the PBMC of a NMI BCa patient **(B)** IFNγ production by total NK cells and **(C)** $CD56^{bright}$ NK cells from BCa patient PBMC or tumor **(D)** LAMP-1 expression by total NK cells with and without Tim-3 blockade (PBMC: n=15, Tumor: n=6). **(E)** Cytolytic effector molecules were measured in PBMC or tumor:K562 co-culture supernatants ± *ex vivo* blockade of Tim-3 with an NK/CD8 cytokine Legendplex kit (BioLegend) (PBMC: n=8, Tumor: n=6). TIGIT blockade (clones MBSA43 or 741170) and TIGIT + Tim-3 blockade were performed as described above. **(F)** Frequency of $IFN\gamma^{+}$ NK cells, **(G)** $IFN\gamma^{+}$ $CD56^{bright}$ NK cell, and **(H)** LAMP-$1^{+}$ NK cells after single or double blockade as indicated s, (TIGIT blockade: PBMC: n=8, Tumo:; n=4), (TIGIT+Tim-3 blockade: PBMC: n=10, Tumor: n=4). Data represent a combination NMI and MI disease. ns, p≥.05, *p<.05, **p <.01, ***p <.001 as determined by paired t-test.

### Figure S4

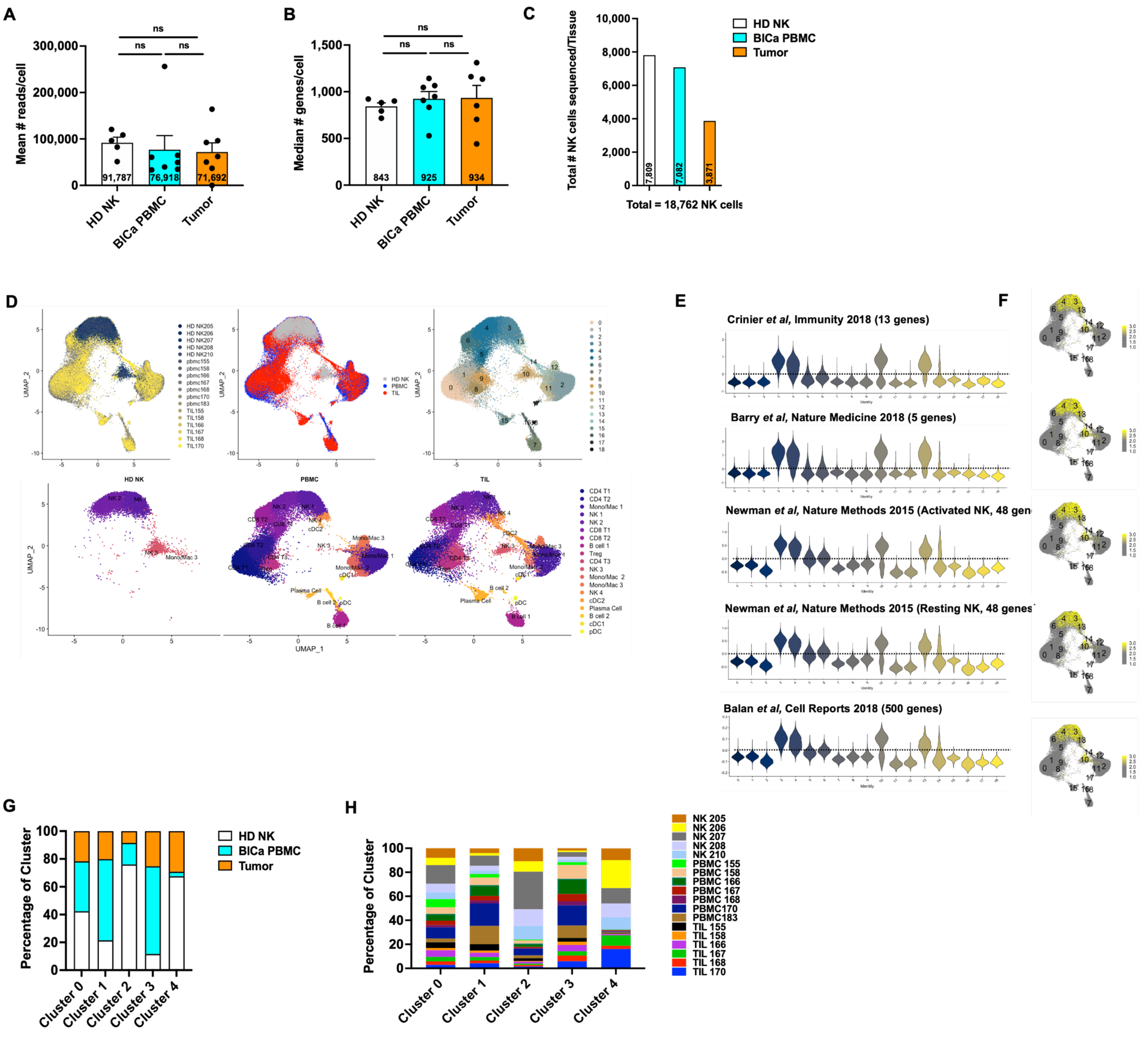
A
Mean # reads/cell
300,000
200,000
100,000
0
ns
ns
ns
91,787
76,918
71,692
HD NK
BlCa PBMC
Tumor
B
Median # genes/cell
1,500
1,000
500
0
ns
ns
ns
843
925
934
HD NK
BlCa PBMC
Tumor
C
Total # NK cells sequenced/Tissue
10,000
8,000
6,000
4,000
2,000
0
7,809
7,082
3,871
Total = 18,762 NK cells
HD NK
BlCa PBMC
Tumor
D
HD NK
PBMC
TIL
UMAP_1
UMAP_2
E
Crinier et al, Immunity 2018 (13 genes)
Barry et al, Nature Medicine 2018 (5 genes)
Newman et al, Nature Methods 2015 (Activated NK, 48 gen
Newman et al, Nature Methods 2015 (Resting NK, 48 genes
Balan et al, Cell Reports 2018 (500 genes)
F
G
Percentage of Cluster
100
80
60
40
20
0
Cluster 0
Cluster 1
Cluster 2
Cluster 3
Cluster 4
HD NK
BlCa PBMC
Tumor
H
Percentage of Cluster
100
80
60
40
20
0
Cluster 0
Cluster 1
Cluster 2
Cluster 3
Cluster 4
NK 205
NK 206
NK 207
NK 208
NK 210
PBMC 155
PBMC 158
PBMC 166
PBMC 167
PBMC 168
PBMC170
PBMC183
TIL 155
TIL 158
TIL 166
TIL 167
TIL 168
TIL 170

**Figure S4: Characteristics of immune cells analyzed by scRNAseq. (A)** Mean number of sequencing reads per cell for the tissues examined. **(B)** Median number of genes detected per cell for each tissue. **(C)** Number of NK cells sequenced per tissue **(D)** UMAP showing immune clusters resulting from the integration of all sequenced PBMC, TIL, and HD NK cells. Cells are colored by donor, tissue, and cluster **(top)** or by annotated cluster split by tissue **(bottom)**. **(E)** Previously reported NK cell gene signatures were used to calculate a module score using the AddModuleScore function in Seurat. Clusters 3, 4, 10 and 13 had the highest NK cell module score for all five signatures tested, and were thus annotated as NK cells and subclustered against each other **(F)** Module scores for previously reported NK cell gene signatures overlaid on UMAP of all immune cells sequenced. Number of NK cells sequenced for each donor. **(G)** Cluster membership by tissue of origin for sub-clustered NK cells. **(H)** Cluster membership by donor for sub-clustered NK cells. Numbers in bars for **A + B** show averages, numbers in **C** show absolute values. Violin plots show the single-cell distribution of NK cell module scores for each of the 19 clusters obtained after aligning 68,192 immune cells from BlCa patient PBMC (n=7),TIL (n=6), and NK cells from HD PBMC (n=5). ns≥.05 *p<.05, **p <.01, ***p <.001, ****p <.0001 as determined by unpaired Mann-Whitney test. Error bars show S.E.M.
